# Plant behaviour: a mathematical approach for understanding intra-plant communication

**DOI:** 10.1101/493999

**Authors:** Fabio Tedone, Emanuela Del Dottore, Barbara Mazzolai, Pierangelo Marcati

## Abstract

Plants are far from being passive organisms being able to exhibit complex behaviours in response to environmental stimuli. How these stimuli are combined, triggered and managed is still an open and complex issue in biology. Mathematical models have helped in understanding some of the pieces in the complexity of intra-plant communication, but a larger and brighter view, setting together multiple key processes, is still missing. This paper proposes a fully coupled system of nonlinear, non-autonomous, discontinuous, ordinary differential equations to describe with accuracy the adapting behaviour and growth of a single plant, by deeply analysing the main stimuli affecting plant behaviour. The proposed model was developed, and here sustained, with the knowledge at the state of the art; and validated with a comparison among numerical results and a wide number of biological data collected from the literature, demonstrating its robustness and reliability. From the proposed analysis it is also shown an emerging self-optimisation of internal resources and feedback stimuli, without the need for defining an optimisation function for the wellness of the plant. The model is ultimately able to highlight the stimulus-signal of the intra-communication in plant, and it can be expanded and adopted as useful tool at the crossroads of disciplines as mathematics, robotics, biology, for instance for validation of biological hypothesis, translation of biological principles into control strategies or resolution of combinatorial problems.

**Author summary:** Plants are able to adapt themselves to a wide range of conditions. The complexity and efficiency of this behaviour suggest a richness of signals exchanged that allowed scientists to talk about communication. Understanding this network of stimuli is an important challenge to increase biological knowledge as well as to adapt plant mechanisms to different research fields. To address this quest we developed a mathematical model able to describe what cues are fundamental in plant communication and how they interact each other.

## 1 Introduction

Plants have developed, over generations, the ability to adapt their resource allocation, foraging strategies and growth rates according to the environmental conditions that change over space and time [1]. Such ability to adjust internal processes and morphology in response to the environmental changes is referred by biologists as plant behaviour [2], or plant plasticity [3].

In a more recent review, [4] redefined the definition of plant plasticity adding to the previous environment-induced stimuli, the ability of plants to live according to purposeful behaviours developed on the base of a sort of memory for some fitness optimisation. Moreover, according to the author, the large number of external stimuli and internal targets that the plant has to manage could be the reason of not placing critical functions in a single (or two) organ, like animals’ heart or brain.

Hence, plants are able to trigger a network of different cues that control molecular signalling in different tissues, organs and the whole plant [5]. Decoupling and clearly understanding such signals are challenging tasks but of fundamental importance for plant behaviour analysis. According to the studies that better analysed plant behaviour [1], intra-plant signals can be categorised into three main groups: I) *electric signals*, typically used by plants to stimulate a fast adaptation, as required for instance in case of some nutrient deficiency or wounds [6, 7]; II) *oscillatory signals*, that are driven by the circadian clock, and direct the growth [8, 9]; and III) *hormons*, like *auxin*, *citokin*, *giberellin*, which are typically messengers of stress conditions [10, 11].

Towards the understanding of plant behaviour, several mathematical models have been proposed with the aim of analysing selected signalling pathways and internal dynamics. For example, in [12], a coupled PDE-ODE (Partial and Ordinary Differential Equation respectively) system describes the cross-talk between the *brassinosteroid* and *gibberellin* signalling pathways; while, other authors modelled the water uptake by roots and the associated morphology [13, 14]. The fluid dynamics inside the xylem and phloem has been also of interest [15, 16], as well as, the photosynthesis [17–20] and the interaction between different plants [21, 22]. However, plant behaviour is determined by a very complex assembling of very different signals and to date none of the proposed models is able to describe such complexity and clearly discriminate stimulus-signal-behaviour chain in view of an internal communication network.

A previous tentative of analysing the mechanisms behind intra-plant communication has been done in [23], with the aim to identify and exploit basic biological principles for the analysis of plant roots behaviour. The analysis was carried in simulated environment and on robotic roots by translating the identified biological rules into algorithmic solutions. Particularly, on each root individually, the uptake kinetic and a local memory were used to adjust the local priority of nutrients, while the resources collected as a whole were shared according to the source-sink principle. Even if each root is acting independently from the others, just considering local information, results shown the emergence of a self-organising behaviour in plant roots aim at optimising the internal equilibrium among nutrients at the whole plant level, with no need for a central coordination or an optimisation function. However, in that work just a small part of the plant complexity was considered, for instance, it completely excluded the analysis of the above ground organs and photosynthesis-related dynamics.

In this paper, we do not aspire having a complete description of the plant complexity, yet we want to identify the main cues influencing the growth of a plant with the aim of investigating on the processes playing a role in the intra-communication for plant growth decisions. We propose and explain here a system of ODEs that, differently from the state of the art models, take into account the entire sequence of processes from nutrients uptake, photosynthesis and energy consumption and redistribution.

In particular, according to [24], biomass is an important estimator of plant fitness but it cannot be studied alone. For this reason, our model focuses on both above and below ground biomass, along with nutrients and sugar levels. In this way, it is possible to study independently leaves and roots adaptations to external and internal conditions. Moreover, unlike the most of existing models, it is possible to describe plant behaviour when a more complete interaction among nutrients is taken into account.

We propose here a system of ODEs to outline the main cues influencing the growth of a plant. In section 2 the model is described in details, emphasising the well proved biological assumptions used to build up the equations. Section 3 presents several results of numerical simulations and their comparison with the biological data obtained from the literature, for validation of the model. Finally, in section 4 are discussed the key elements of our model, its functionalities and the improvements with respect to the state of the art, while section 5 presents conclusive remarks.

## 2 Methods

### 2.1 Description of the mathematical model

Through the photosynthesis, plants produce glucose from carbon dioxide and water, using the energy from sunlight. To avoid starvation in the night, they should carefully manage the resources acquired and produced during the day. Also, a plant is composed by organs with different tasks. For instance, roots are the organs of the plant furnishing water and nutrients to the above ground part, receiving from it the photosynthetic products, particularly in the form of sugars, needed for growth and maintenance. While the leaves are the energy producers and starch stores used during nights. According to the resources availability and the energy produced, the biomass is differently distributed.

Nutrients are transported toward root surface mainly by water flow in soil. Nevertheless, nutrient uptake is controlled by spatial distribution of roots. In fact, as explained in [25], root system can develop to easily access to immobile nutrients (e.g. phosphorus or ammonium), or locally adapt their exploitation power for a specific nutrient.

According to [26], there are 17 most important nutrients for plants that can be distinguished in two main categories: macronutrients (usually required in high quantities) and micronutrients (which are required in smaller quantity).

As reviewed in [27], the nutrients content in soil should fall in specific ranges for the plant not to suffer from toxicity or deficiency effects. Moreover, the same authors show how an optimal ratio between nutrient contents in plant is necessary and how an imbalance of such ratio can actively affect the uptake of nutrients.

Due to the complex network of interactions between nutrients, in both soil and plant, usually only one nutrient [17, 20, 28], or the single water availability [14, 21, 29], have been studied.

Some authors have studied an interaction between nutrients but avoiding a whole plant interaction like photosynthesis or sugar-signalling [30].

In our model, in order to study a more complete network of signals, over sugar-signalling, we developed a root process dynamic model considering the presence of two nutrients in the soil, and assuming a non-limiting and uniform distribution of water in soil and of CO_2_ in air.

For our study, we selected nitrogen (n) and phosphorus (p), being both fundamental nutrients for growth in plants.

Nitrogen is the most abundant nutrient in plants, essential constituent of protein and chlorophyll. Moreover it promotes root growth, improves fruit quality, enhances the growth of leafy vegetables, increases protein content of fodder, encourages the uptake and utilisation of other nutrients [31].

Phosphorus is essential for growth, cell division, root lengthening, seed and fruit development, and early ripening. It is a part of several compounds including oils and amino acids. The P compounds adenosine diphosphate (ADP) and adenosine triphosphate (ATP) act as energy carriers within the plant [32].

The proposed model takes in consideration all these fundamental working principles, by describing the dynamics of each actor: photosynthesis, starch, sucrose, nutrients uptake and management, costs of maintenance and growth, and biomass allocation.

For the sake of simplicity, the time dependence of functions and variables is implied and it will not be reported in the following model equations.

### 2.2 Photosynthesis

We described the photosynthesis 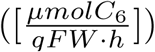 process with a time-dependent function describing the content of sugar produced by each gram of photosynthetically active leaf biomass:

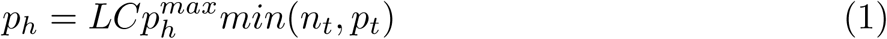

where:

- *L* [−] is a binary function to distinguish day by night. It is 1 during day and 0 during night. It should be noted that L can be easily modified in a continuous time-dependent function assuming values in the range [0, 1], so to consider temperature and light intensity effects.
- *C* [−] is a positive function in the range [0, 1]. It represents a limiting coefficient to photosynthesis dependent on the plant status. In fact, different saturating processes can affect stomatal conductance, reducing photosynthesis production ([33] reviewed many modelling approaches). Particularly, [34] suggests a more specific photosynthesis limiting process, which is indirectly linked to the need of the plant of avoiding the production of starch and sugar in excess. Accordingly to [35] and [20], we propose a limiting function depending on the maximum starch that the plant can consume in nightly hours. The C function can be written as:

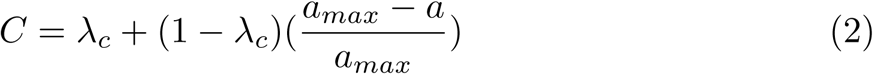

where:

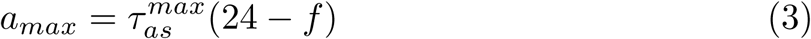 The parameter *λ*_*c*_ describes the strength of photosynthesis limitation, a is the starch, *a*_*max*_ the maximum starch that can be consumed at the maximum rate 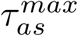 during the night and f the photo-period. The sugar limitation will affect the starch partition (*γ*) dynamics, that will be explained later (Section 2.3.2).
- 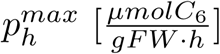 is the maximum rate of photosynthesis. As for in [36], it is fixed to 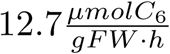.
- *n*_*t*_ and *p*_*t*_ represent nitrogen and phosphorus saturating threshold, respectively. Both these nutrients play a role in the photosynthesis process (see [37] for an explanation of the role of nitrogen, while [38] reviews phosphorus influence on photosynthesis). Hence, photosynthesis is strictly correlated with nutrient availability in a saturating way. In fact, according to the law of minimum [39], the most limiting nutrient content will mainly affect the photosynthesis. By [17], the rate of carbon production obtained from nitrogen content is estimated as:

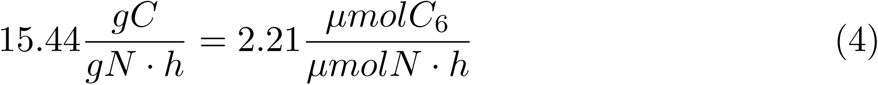 Hence, the minimum nitrogen content required to sustain the maximum rate of photosynthesis is:

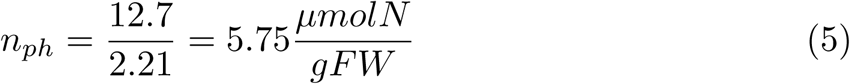 The saturating function *n*_*t*_ can be computed as:

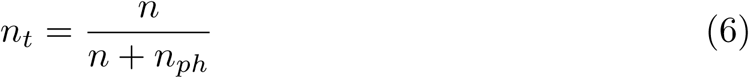 Not having found similar information for phosphorus, we estimated the saturating function of this nutrient by assuming the consumption of phosphorus as one-tenth of the nitrogen’s one, according to the optimal ratio among these two nutrients, which has been observed to be about 10 (as reviewed in [23]). Therefore, we approximated:

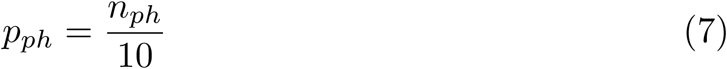

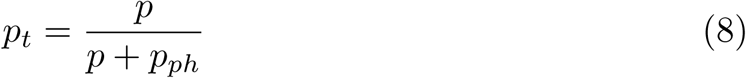

where 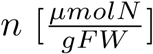 and 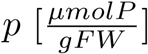 are respectively the nitrogen and phosphorus content.

In particular, if *n* < *n*_*ph*_ or *p* < *p*_*ph*_, then it should be *n*_*t*_ = 0 or *p*_*t*_ = 0 respectively just because there are not enough resources to start the chemical processes.

It should be noted that Eq (1) is a simplification of the more general formula:

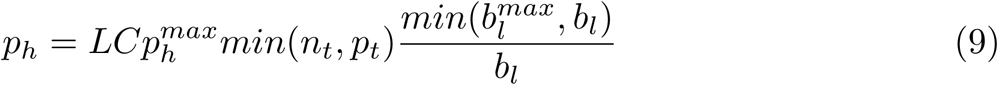

which takes into account the aboveground biomass (*b*_*l*_) and the maximum photosynthetically active leaf biomass 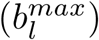. However, since in all papers used for validation there was no clue that the critical biomass was reached during the experiments, the 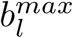 is assumed to be infinite, allowing us to remove the biomass factorial term.

### 2.3 Starch and Sucrose

During the day, some photosynthate is stored as starch to be degraded during night to sustain the nocturnal metabolism. Independently by the length of the night, the starch is degraded into sucrose in a nearly linear manner (at a rate *τ*_*as*_), such that almost all of the starch is used by dawn [40]. We described starch and sucrose dynamics with the equations:

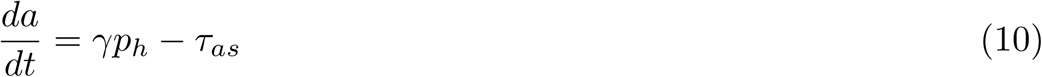

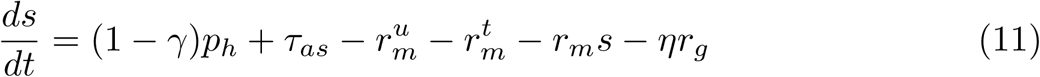

A very similar dynamic can be found in [41] and [42]. Nevertheless, some important differences deserved to be outlined:

- The photosynthesis *p*_*h*_ in the current model is not a constant, but it is a time-dependent function parametrized on nutrient availability, light, sugar-signalling and leaf competition (as described in previous section);
- The starch partition coefficient *γ* is not a constant, neither it is estimated assuming sucrose homoeostasis (as in the model of [43]). It provides instead plant adaptation as a function of time (its dynamic is described in Section 2.3.2);
- The starch degradation rate *τ*_*as*_ is simpler respect to the definition provided in [41] and [42], and it does not depend on a subjective dusk [44]. Even considering the simplification adopted in this paper for starch degradation, its dynamic remains correct (see file S1 in supporting information for demonstration);
- Just like in [20], both uptake 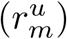 and transport 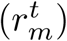 processes are sucrose consuming, over the usual maintenance respiration (*r*_*m*_);
- The rate of sucrose sent into the phloem intended for growth (*ηr*_*g*_) is affected by nocturnal efficiency [45].

#### 2.3.1 Starch degradation

It is well established that maintenance and growth respiration occur both during day and night. The sucrose to sustain nightly metabolism is provided by the degradation 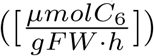 of starch stored in the morning [40], and here defined as:

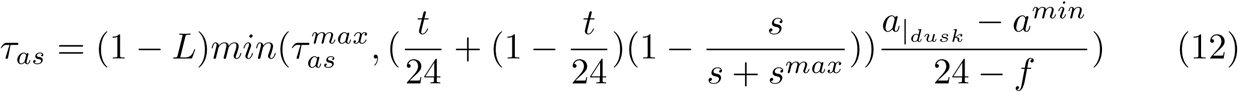

where:

- 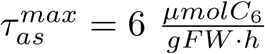 is the maximum degradation rate estimated by the results shown in [36];
- 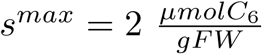 is the maximum sucrose content in leaves. It is estimated by experimental values in [46]. A negative feedback on starch degradation due to high sucrose levels is proposed also in [47]. Hence, the introduction of a maximum threshold is justified by these evidences. On the other hand, a minimum threshold should be expected to trigger sugar production or starch degradation in case of starvation. Like in [36], *s*^*min*^ is estimated as 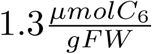;
- 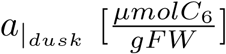 is the starch content at the beginning of the night. The use of this parameter makes us highlight some differences with respect to the model presented in [41], where: I) the function proposed to describe starch degradation is independent from this parameter and has a peak at dawn; II) starch degradation is assumed to exist also during the light period; and III) the non linear discontinuous function proposed takes into account the surface of starch granule. In the supporting material, we integrated their function in our model and we compared the two approaches;
- 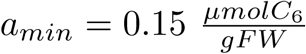 is the minimum amount of starch that plant has to ensure. It is estimated by [41].

Hence, during night (*L* = 0), the starch is degraded with a rate that depends both on the time t with respect to the next expected dawn (circadian clock), the sucrose signalling and the constant rate necessary to consume almost all of the starch by dawn. This strategy holds unless the required rate is greater than the maximum one.

#### 2.3.2 Starch and sucrose partitioning

[47] explains how rising levels of sucrose stimulate starch synthesis, while daily sucrose starvation decreases starch accumulation. Nightly sucrose starvation, on the other hand, promotes starch production [20]. From these observations, we defined the dynamic of starch-sucrose partitioning as:

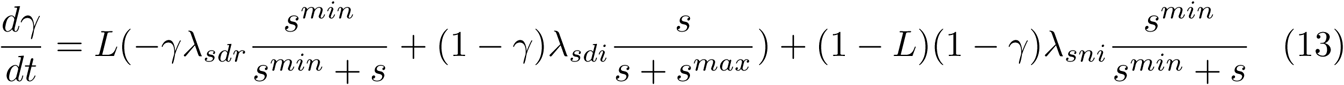

### 2.4 Biomass allocation

Sucrose transport is essential to allow growth and maintenance of non photosynthetically active tissues like roots, shaded and/or young leaves, and other storage organs [48]. Actually, the sucrose is loaded into phloem generating an osmotic pressure which drives both water and sucrose from source tissues to sinks [49].

In particular, testing Münch hypothesis, [50] shows a sink priority concept affecting how sucrose loaded is allocated between sinks. In our model, sucrose is divided between above and below ground biomass. The partitioning coefficient *f*_*r*_ of sucrose allocated to roots is described by the following equation:

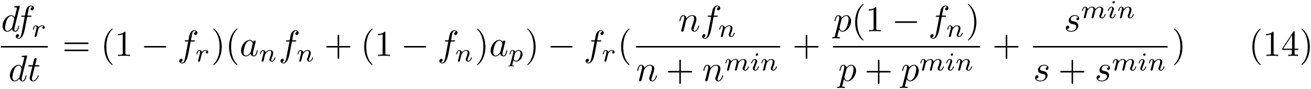

where 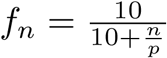 is a stoichiometry signal with respect to the optimal nutrient ratio. The frequency parameter fixed to 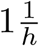 is omitted in the previous equation.

According to [51], the less nutrients are, the more root priority is and vice versa. In particular, [52] stresses that sink priority is affected by the most limiting resource, while [53] and [54] highlight how plants are able to adapt their allocation strategies according to internal and external stimuli. Hence, while functions *a*_*n*_ and *a*_*p*_ describe nutrients demand, the value of s with respect to the threshold *s*^*min*^ describes a limitation due to leaves.

As result, the function *f*_*r*_ balances the nutrients demand with respect to the most limiting resource, and describes the sucrose allocation between leaves and roots. Moreover, being a central brain missing in plant, there is not central control on carbohydrates allocation, thus we hypothesis a different nutrient feedback driving resources allocation within roots (the same hypothesis should be valid for leaves, but we are not interested in modelling spatial distribution of leaves at this stage of the work).

According to [55], [51] and [56], the innate soil heterogeneity affects root system morphology promoting root proliferation in nutrient-rich soils. In our model the choice is between two different soil zones. Given sucrose allocated to roots, the proportion of resources sustaining growth in the top soil is given by the following weighted average ([−])

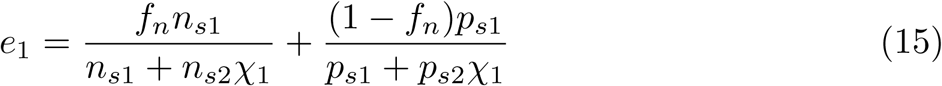

Being soil zones vertically distributed, the binary function *χ*_1_ is necessary to express if the root biomass has reached the second (deeper) soil zone. Actually, *χ*_1_ is 1 if 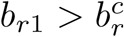, where 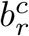 is a parameter describing the critical biomass to reach before having access to the deeper zone.

### 2.5 Growth

Considering the above observations on biomass allocation, we can describe the growth dynamics of leaves and roots with the equations:

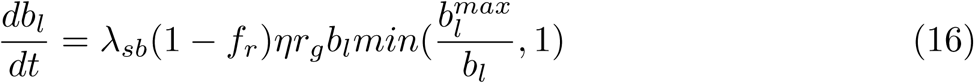

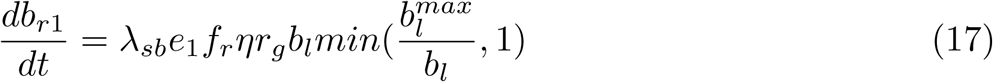

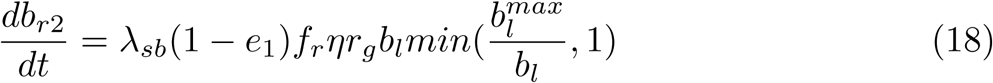

In Eq (16), (17) and (18), *λ*_*sb*_ is a conversion parameter from sucrose to biomass. The growth, according to the specific tissue priority (*f*_*r*_ and *e*_1_), will be proportional to the whole sucrose exported by photosynthetically active leaf biomass.

### 2.6 Nutrients dynamics

To study the root system adaptation in a changing environment, we assumed the soil divided into two zones: the topsoil and the subsoil. This choice is justified by the different mobility of phosphorus and nitrogen (the selected nutrients for our study). The former is immobile and usually a greater content can be found in topsoil [57]. Nitrogen, on the other hand, is quite mobile and easily transported by percolating water in subsoil [58, 59].

We described the plant dynamics of nitrogen 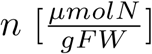 and phosphorus 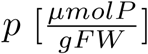 content as:

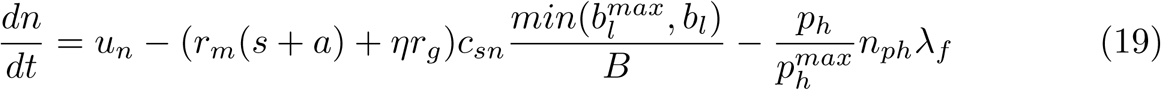

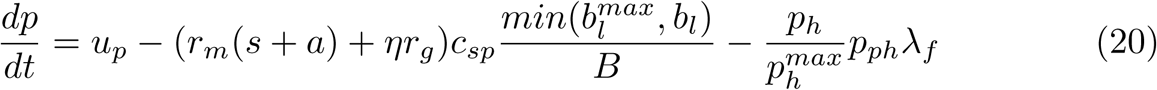

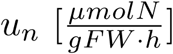 and 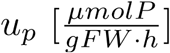 are the nitrogen and phosphorus uptake rate, respectively (see Section 2.6.1 for details).

In [17], the cost of assimilating *n* into biomass is estimated as 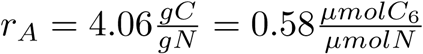. Hence the *n* cost of assimilating C_6_ can be estimated as proportional to:

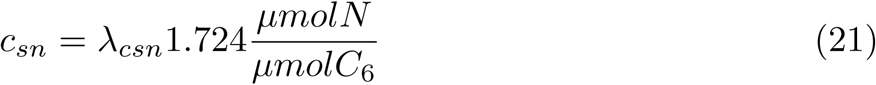

As already previously justified (Section 2.2), the *p* cost of assimilating C_6_ can be approximated using the optimal ratio value ([23]) as 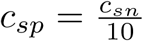. In particular, metabolism, transport and growth will be reduced if the nutrient content is not sufficient to sustain the costs (*c*_*sn*_ and *c*_*sp*_). Then:

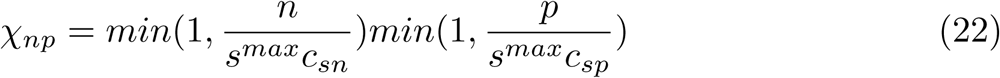

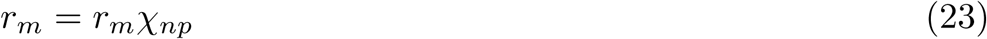

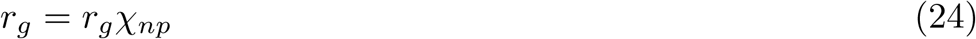

The second terms of Eq (19) and (20) are the *n* and *p* costs owed to the use of sucrose for maintenance and growth. Also, the starch requires a maintenance respiration (and then a cost in nutrients) due to the production of storage structures [60].

Note that s and a represent the quantity of moles of C_6_ for each gram of photosynthetically active leaf biomass. Then, the term 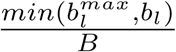 is necessary to consider the actual plant sugar content, being B the full plant biomass.

Finally, the last terms of Eq (19) and (20) express the nutrient costs due to photosynthesis. Knowing the nutrient cost for the maximum rate, *n*_*ph*_ (Eq (5)) and *p*_*ph*_ (Eq (7)), the actual consume will be proportional to the intensity of photosynthesis. The frequency parameter 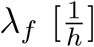 will be estimated later (Section 2.9).

#### 2.6.1 Nutrients uptake

Having nutrients located in two discrete soil zones, we described the potential uptake with two terms (labelled in subscript with index 1 - topsoil - and 2 - subsoil), each multiplied by a factor that takes into account the whole biomass. For nitrogen (and similarly for phosphorus) we defined:

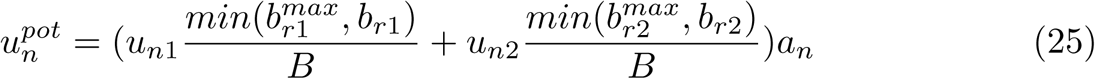

where *b*_*r*1_ is the root biomass in topsoil, *b*_*r*2_ the root biomass in subsoil. 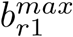and 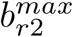 are the maximum root biomass allowed in each soil zone. As for 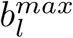 (in Eq (9)), we fix to infinite their value, assuming no root competition. *a*_*n*_ is an internal feedback explained later.

A limiting factor [−] should be taken into account. Its role is to decrease the forage if the hourly potential uptake 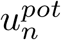 needs excessive sugar consumption with respect to the available one. Thus:

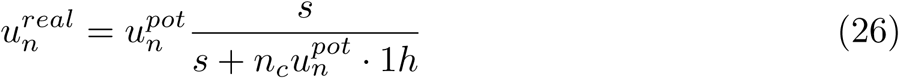

*u*_*n*1_ and 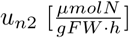 are the nutrient rates of uptake, while *n*_*c*_ will be estimated later. It is well established that the uptake follows a Michaelis-Menten kinetics [61, 62]:

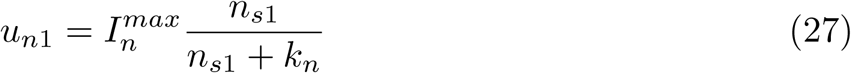

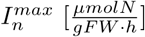 is the nutrient maximum uptake, 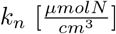 is the Michaelis Menten constant and 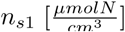 is a time depending function describing the nutrient content in the soil zone. Similar equations are defined for *u*_*n*2_ and phosphorus in top and subsoil.

Several papers have shown how Michaelis Menten parameters are not fixed [63–65]. Plants, in fact, are able to modify these parameters to be more or less affine with the specific nutrient. This adaptive behaviour is experimentally measured changing nutrient soil content [66].

However, [55] notes how changes in kinetics parameters could be due to plant internal status instead of environmental reasons. In particular, [55] reports how root proliferation and uptake require varying energies, and how the balance between costs and gains could explains the plant strategy.

In our model we tried to describe this behaviour. For this reason, from [67] we estimated kinetic parameters for nitrate, and from [63] we approximates phosphorus parameters:

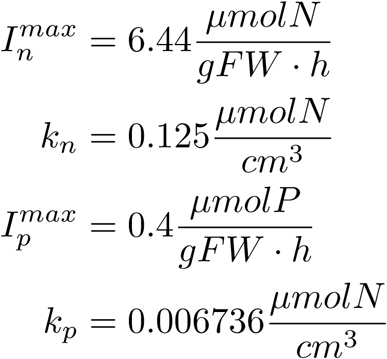

They represent the maximum values for the couple (*I*^*max*^, *k*). To simulate the nutrient affinity adaptation we introduced in Eq (25) the internal control *a*_*n*_ [−]. An analogous control *a*_*p*_ is inserted for phosphorus dynamics

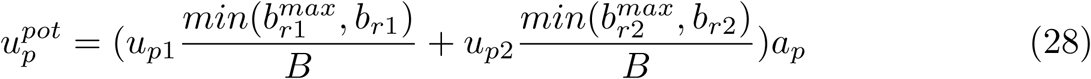

Both *a*_*p*_ and *a*_*n*_ are positive time dependent functions bounded by 1. Their dynamics are described below (again the frequency parameter fixed to 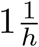 is omitted in both the following equations):

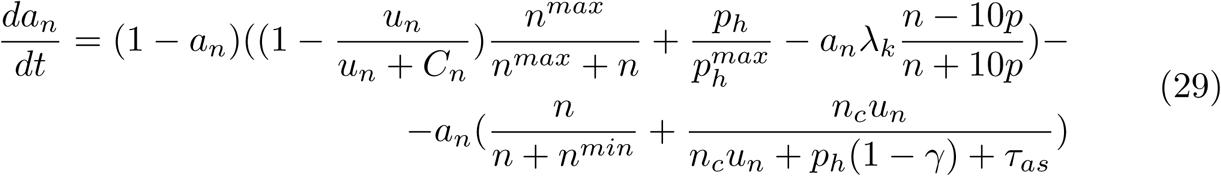

A similar formulation holds for the phosphorus control function *a*_*p*_:

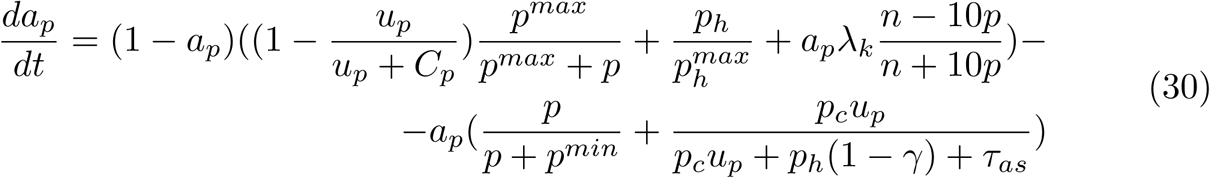

*C*_*n*_ and *C*_*p*_ represent the nitrogen and phosphorus costs due to the actual strategy. They are the absolute value of the sum of the second and third terms in Eq (19) and (20).

*n*^*max*^ and *p*^*max*^ take into account the memory of a plant. In fact, according to [18], plants manage its foraging strategies in order to avoid both scarcity and excess of nutrients in stores. In particular, the optimal nutrient status is defined by how many days plant can survive maintaining the same rate of growth if a specific nutrient is not available anymore. [18] estimated for herbaceous plants (like *Arabidopsis*) a memory of 4 days. We fixed *D* = 4 [−]. Hence, the minimum amount of nitrogen content 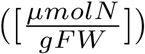 required to sustain photosynthesis and respiration for one day at the maximum sugar content is:

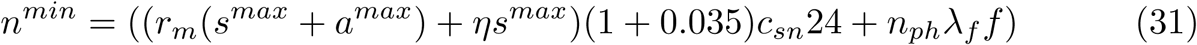

Then, we estimated 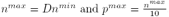. The term:

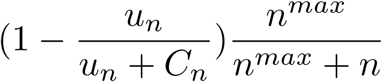

describes the active incentive for the uptake rate according to the internal status and the cost-gain balance as reported in [55]. On the other hand, the last terms in Eq (29) and (30) represent the active limitation to the uptake due to the same concepts.

Moreover, as emphasised by [68], over an active control of nutrient affinity, the uptake rate is passively increased by photosynthesis and leaves transpiration. The term:

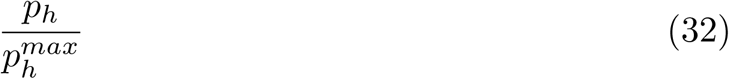

describes this behaviour.

### 2.7 Costs

All the sucrose s produced by photosynthesis and starch degradation is used for maintenance and growth respiration costs.

As in [42], we assumed the respiration to be proportional to the sucrose content. The former has a frequency of 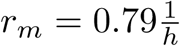. The frequency of sucrose loaded into phloem for growth has a frequency of 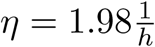. However, as observed in [45], the nightly starvation reduces the growth rate. To take into account this negative feedback, we reduced *r*_*g*_ under the parameter *γ* that in the model describes the nightly efficiency of starch. Hence:

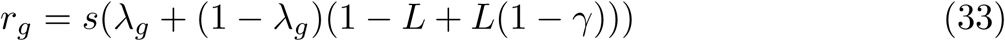

The more *γ* increases, the more *r*_*g*_ decreases. Over maintenance and growth respiration, sucrose is consumed for the transport of carbon along the plant and for the uptake of nutrients. [20] estimates costs of uptaking phosphorus as:

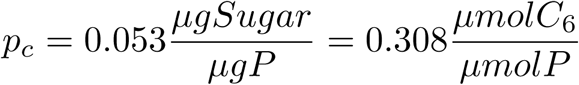

with a sugar transport cost of 0.035 [−] (carbon consumed for each gram of carbon allocated in phloem). The cost of uptaking nitrogen (*n*_*c*_) is assumed, in our model, to be the same of phosphorus (*p*_*c*_), being the uptake mechanism similar among all chemical elements. Hence:

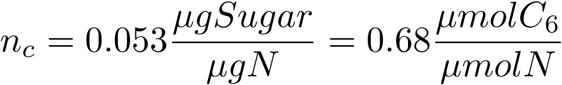

Named 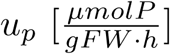 and 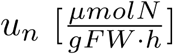 the phosphorus and nitrogen rates of uptake respectively, the whole uptake 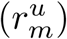 and transport 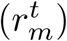 costs are:

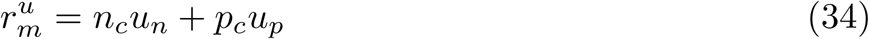

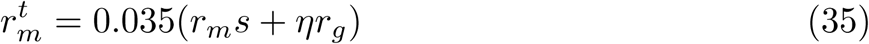

### 2.8 Further environmental contributes

Additionally, we should consider how deficiency or excess of nutrients in soil affect plant behaviour. For instance, [69] shows how in nutrient-poor soils, the secondary metabolism and the plant defence overcome the growth stimulus. For this reason, it is important to quantify nutrient excess or deficiency with respect to an optimal threshold. In [70] for free N medium the value of 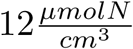 is used, and this value is used as optimal threshold in our model for nitrogen. [66] defined treatments in low P content up to 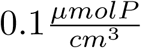, while treatments in high P content start from 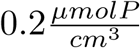. Hence, we fixed the optimal value for phosphorus to 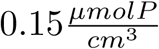. Therefore, the excess or deficiency will be measured with respect to the distance from the defined optimal thresholds and the amount of relative roots that are experiencing it:

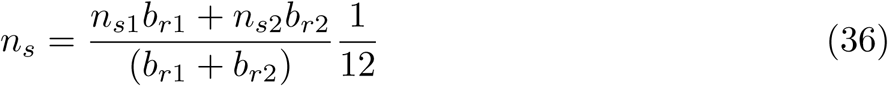

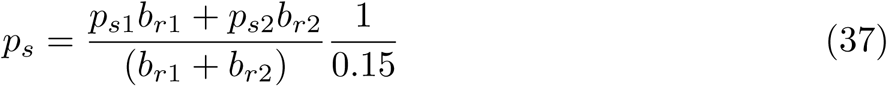

The increased defence-signal can be simulated limiting the conversion parameter *λ*_*sb*_, and the increased metabolism will lead to greater nutrient stocks. We can simulate it increasing *n*^*min*^ and *p*^*min*^. Both N and P are primary for plant growth and a negative feedback should be present to consider their combined effect.

The function *R*^−^ (defined in Eq (38)) is used to reduce *λ*_*sb*_, regulating this way toxicity conditions, and it is symmetric with respect to *n*_*s*_ and *p*_*s*_, with its maximum in (*n*_*s*_, *p*_*s*_) = (1, 1). Moreover it is null if both nitrogen and phosphorus are missing in soil ((*n*_*s*_, *p*_*s*_) = (0, 0)), while all negative values are also cut to 0.

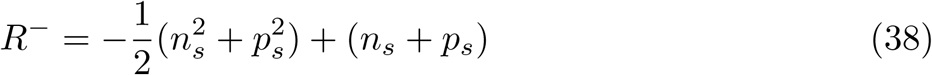

Hence:

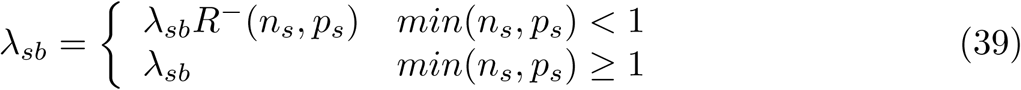

The function *R*^+^is instead used to increase nutrient demand, to regulate deficiency conditions in soil, and it is again symmetric and positive. It has one minimum *R*^+^(1, 1) = 1 and an arbitrary value was fixed for *R*^+^(0, 0) = 139. Hence:

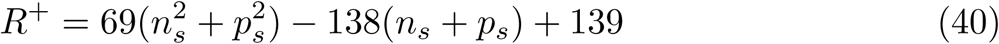

and

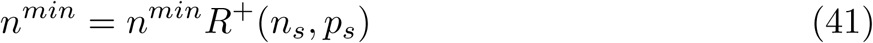

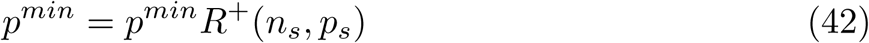

where *R*^+^(*n*_*s*_, *p*_*s*_) = 1 if *min*(*n*_*s*_, *p*_*s*_) ≥ 1.

**Fig 1.**
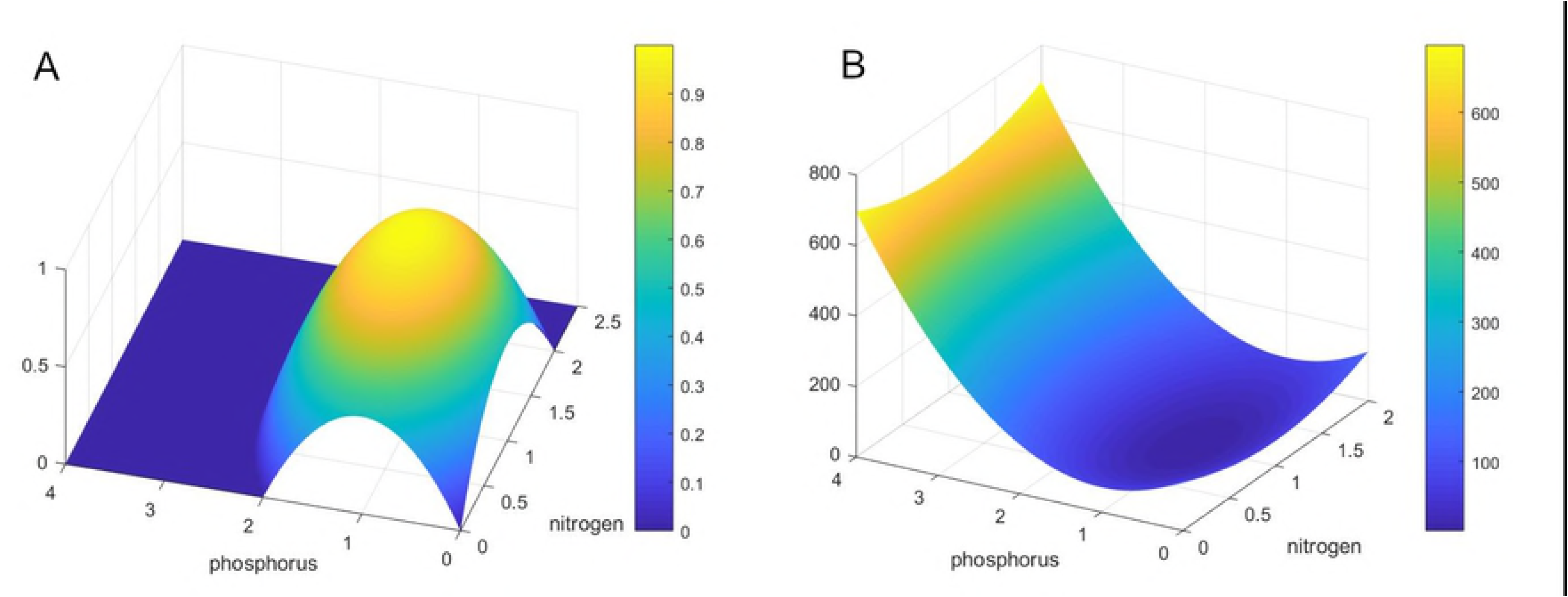
Reduction functions. (A) Example of *R*^−^ function nitrogen or phosphorus soil content is less than the optimal one. (B) Example of *R*^+^ function when nitrogen or phosphorus are deficient in soil.

Going over toxicity levels of nutrients, it will induce negative feedback in plant internal processes. For instance, excess of some forms of nitrogen can change soil pH making phosphorus more available [71]. Other forms of nitrogen can modify the C:N soil stoichiometry and stimulate a greater demand of other micro-nutrients (with possible negative effects) [71]. Moreover, the high nitrogen content in plant can increase the secondary metabolism instead of growth [69]. Other examples come for the phosphorus. According to [71], an high phosphorus content in soil can promote the luxury uptake of P. The plant absorbs more P than it needs modifying P:Fe and P:Zn stoichiometry ratios in plant. Consequences are the same of a Fe and Zn deficiency.

As case study, we wanted to include in our model the plant behaviour in case of luxury levels of P. To this aim, we should insert new equations for Fe and Zn dynamics and consequently more parameters to estimate optimal stoichiometry ratio. However, these data are not easily available in the literature. Hence, we introduced in the model a function simulating Fe deficiency when P content overcomes an optimal threshold. Iron is fundamental in many metabolic pathways and photosynthetic organs. In fact, [72] shows how Fe deficiency affects the development of chlorophyll, limiting the rate of photosynthesis. In the paper, the authors conduct some experiments pointing out the strength of this inhibition. It can induce up to a 50% decrement in chlorophyll accumulation. Moreover, [72] explains that is not possible to induce a greater decrement. Hence, if *p*_*s*_ ≥ 1, the maximum photosynthesis is reduced depending on the amount of phosphorus excess, according to the following:

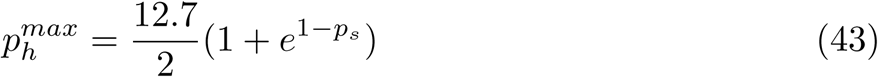

If there is no excess, the photosynthesis does not experience any limitation 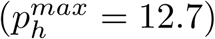. The more excess, the more reduction occurs till the 50%.

### 2.9 Model overview and parameters

Summarising, the model is composed by 11 non-linear equations:

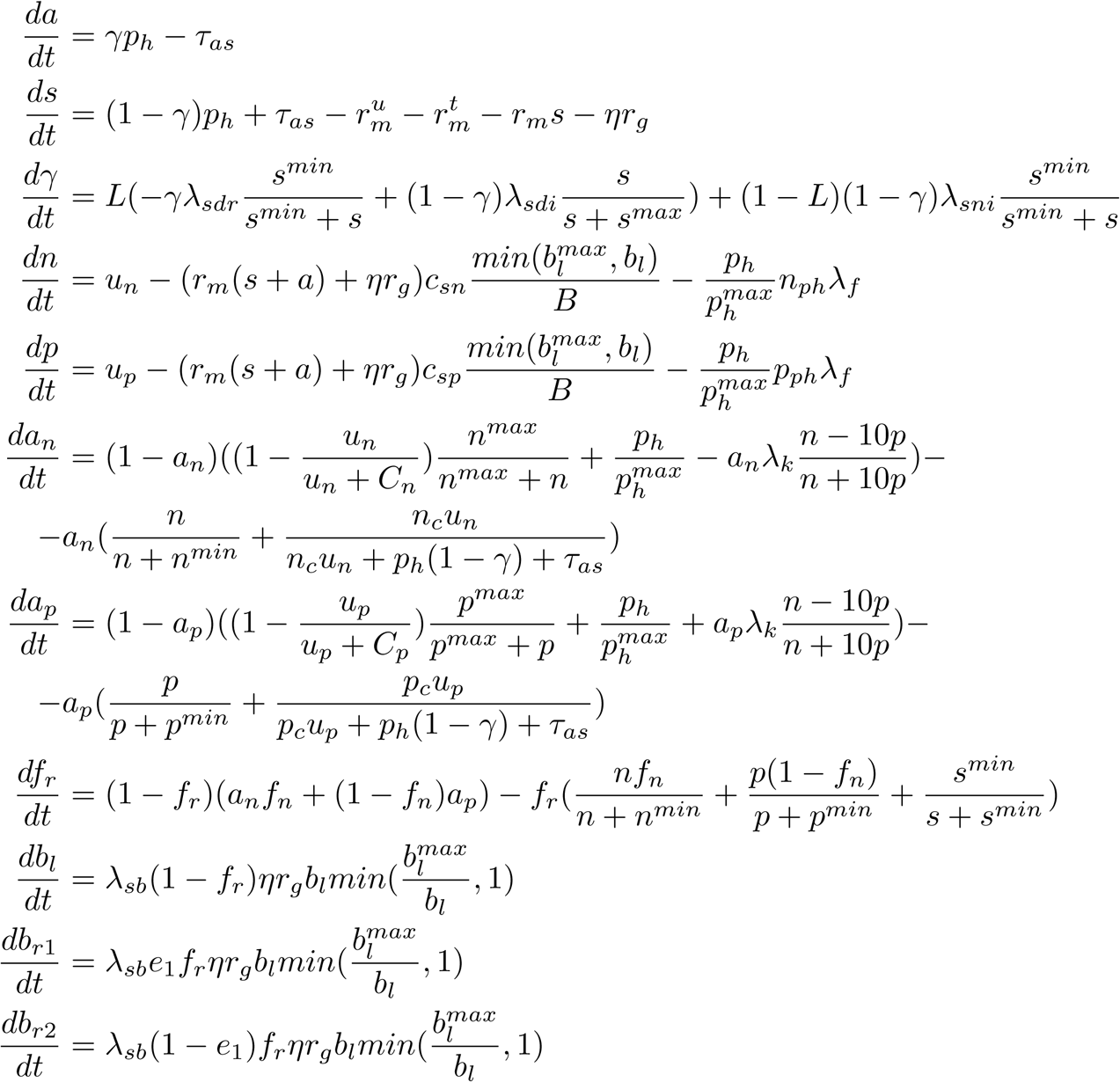

Many model coefficients have been estimated by the literature, as already deeply described in the sections above. The remaining free parameters are:

- *λ*_*c*_ [−]: related to the strength of starch accumulation on photosynthesis (Eq (2))
- 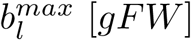: maximum photosynthetically active leaf biomass (section 2.2)
- 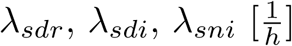: parameters for *γ* dynamics (Eq (13))
- *λ*_*g*_ [−]: related to nightly starvation signal on growth (section 2.7)
- 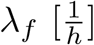: frequency parameter in nutrient consumption during photosynthesis (Eq (19) and (20))
- 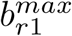and 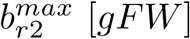: maximum root biomass limiting uptake in each soil zone (section 2.6.1)
- *λ*_*k*_ [−]: coefficient related to the strength of stoichiometry signal in controlling uptake (Eq (29) and (30))
- 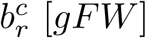: critical root biomass to reach subsoil zone from topsoil (section 2.4)
- 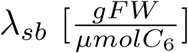: conversion parameter from sucrose to biomass (Eq (16), (17) and (18))
- *λ*_*csn*_ [−]: proportional parameter for *c*_*sn*_ (section 2.6)

In order to estimate them, we used the data available in [46]. In this work, the authors grew *Arabidopsis* under 5 different photoperiods (4*h*, 6*h*, 8*h*, 12*h*, 18*h* of light) for 29 days assuming not limiting medium growth. Moreover, they measured sucrose and starch content and the aboveground rate of growth.

Within our model, we assumed only one soil zone (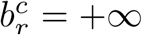 and 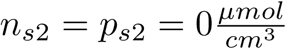). Moreover, as already said, no competition effects were taken into account 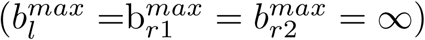.

Of course, in estimating parameters we have to consider a strong dependence of natural processes by temperature. Since the influence of temperature was not explicitly modelled, we estimated different parameters for each photoperiod from [46], using a spline interpolation to approximate different day-lengths. For sake of simplicity, we tried to keep fixed the most of parameters. The best fitting was obtained assuming as variables only *λ*_*c*_, *λ*_*sni*_, *λ*_*sb*_, obtaining the following values:

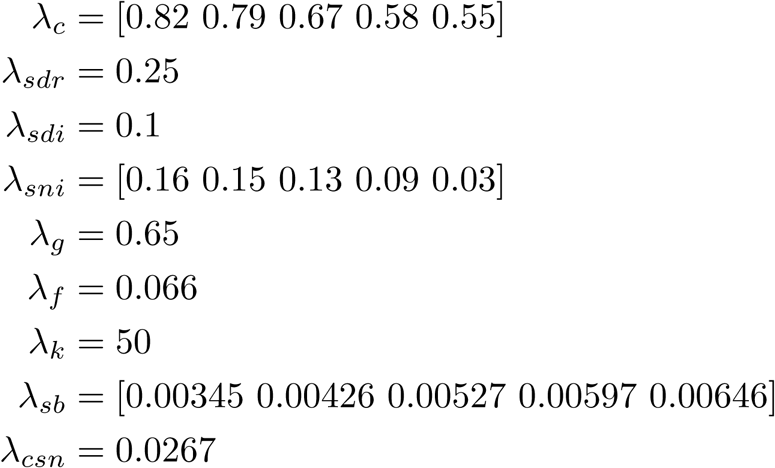

All experiments available to fit and validate the model are conducted in a vegetative phase and it was no possible to estimate a tissue death coefficient. Then, to avoid a greater number of free parameters, no explicit death is accounted in the model, even if the generalisation will be straightforward when new experiments will be available. However, a death concept can be considered present in the model by considering null the growth of the tissues (results for this consideration are reported in section 3.3).

## 3 Results

### 3.1 Model validation

We compared the results obtained by our model with the results extracted from the biological data available in [46].

We looked at the starch dynamic over different lengths of the day: 4, 6, 8, 12 and 18 hours (Fig 2A-C and Fig 3A-B). In each plot, red lines are piece-wise linear functions interpolating data from [46], while green lines are the numerical results obtained from our model. Blue dots represent the dawn (beginning of the day) and red ones the dusk (beginning of the night). The starch grows during the period of sunlight, and decrease during night with a good match between numerical and biological results.

**Fig 2.**
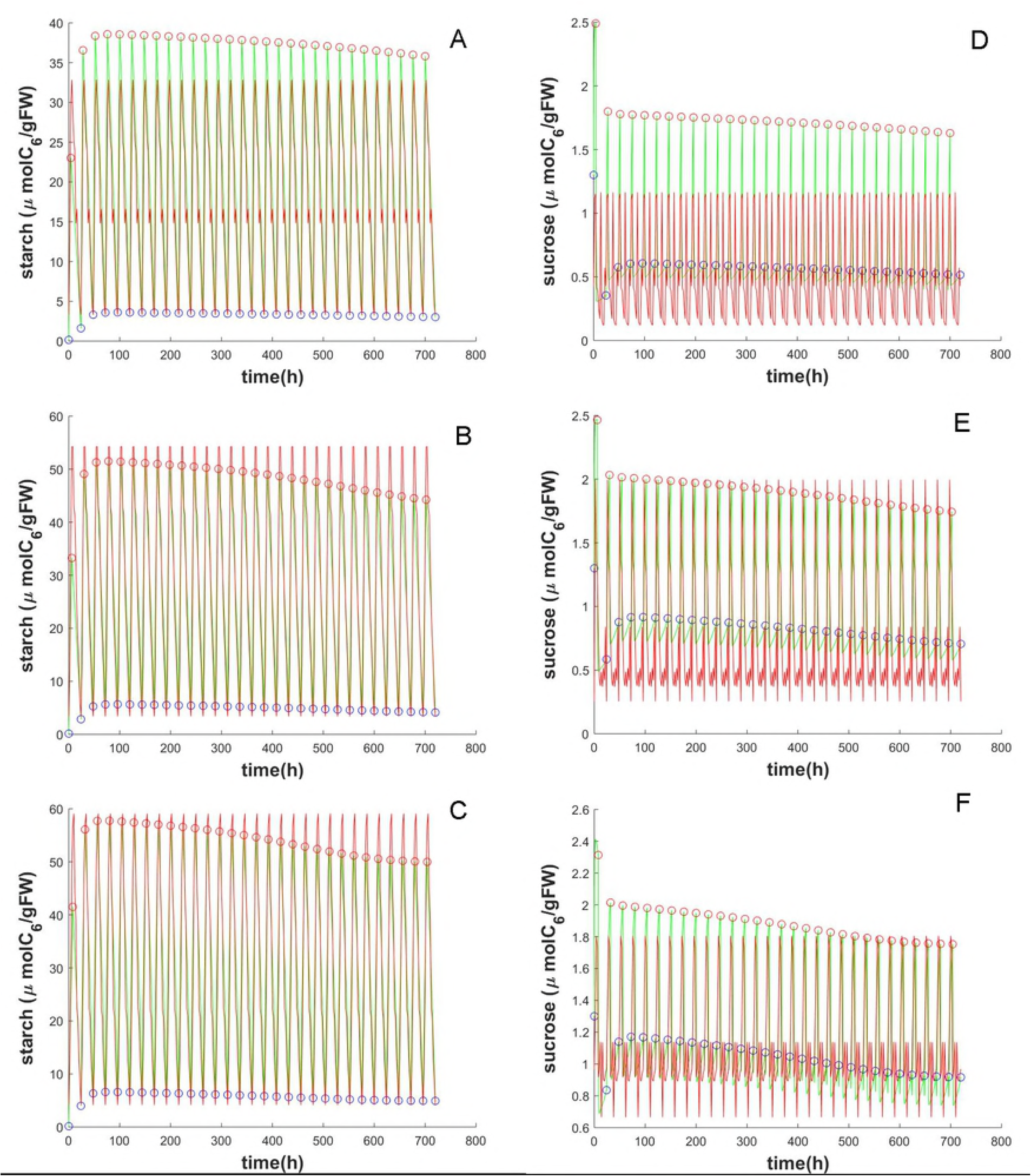
Validation test in short days. Starch (from A to C) and sucrose (from D to F) dynamics for (A, D) 4h day length, (B, E) 6h day length, (C, F) 8h day length. The red lines represent the data obtained from [46], green lines are obtained from our numerical simulations. Blue dots are the beginning of the day and red dots are the beginning of the night.

**Fig 3.**
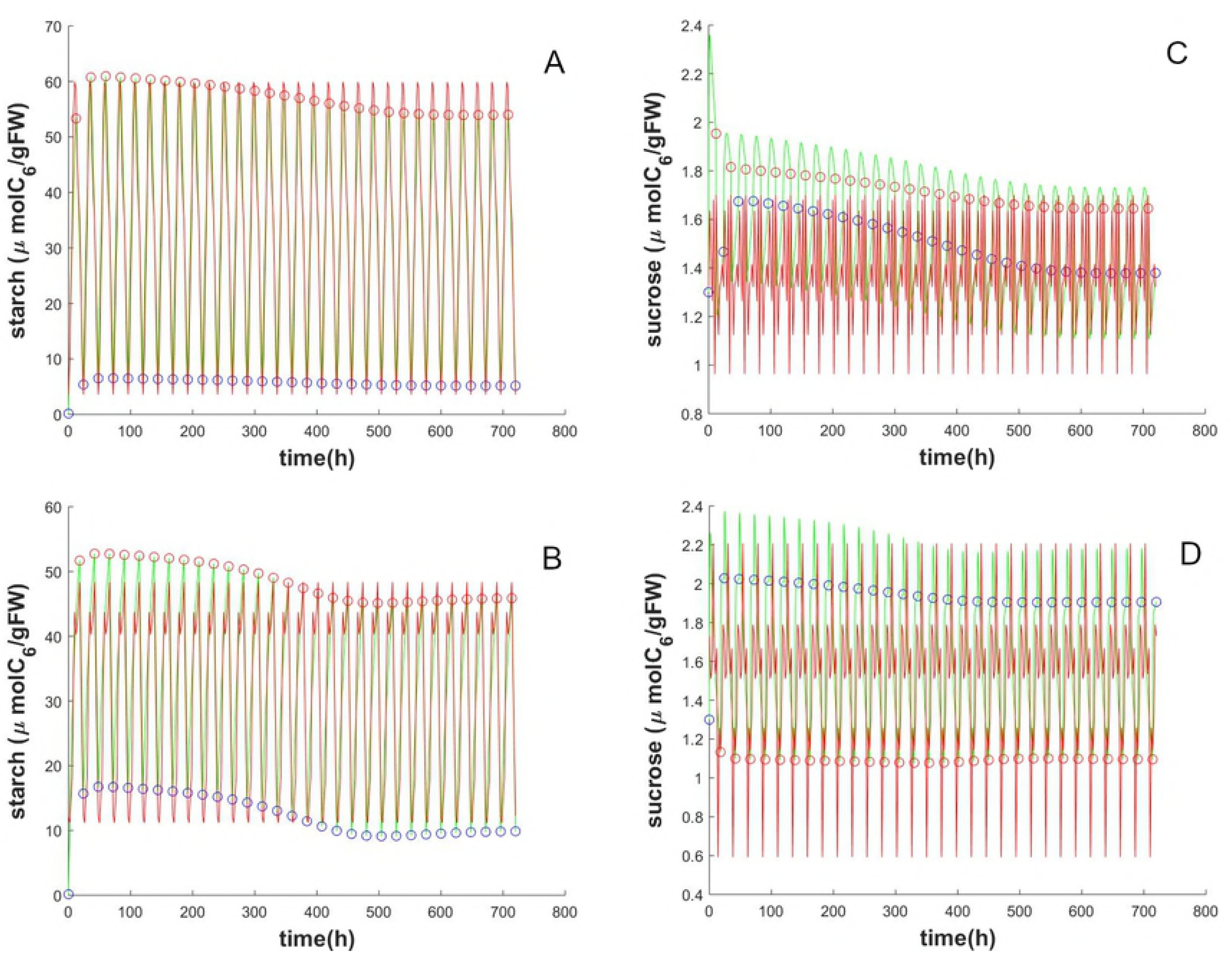
Validation test in long days. Starch (in A and B) and sucrose (in C and D) dynamics for (A, C) 12h day length, (B, D) 8h day length. The red lines represent the data obtained from [46], green lines are obtained from our numerical simulations. Blue dots are the beginning of the day and red dots are the beginning of the night.

The behaviour of the sucrose from numerical simulation (Fig 2D-F and Fig 3C-D) can be considered in average in agreement with biological data, even though less match is found between peaks of dawn and dusk. In fact, the oscillations in sucrose content are smoothed in the model. For instance, we can notice in 4h photoperiod (Fig 2D), instead of having two sucrose peaks during night, the model is able to simulate only one higher peak. A smoother behaviour is instead more evident in 18h of light (Fig 3D); a closer view on the oscillations for this photoperiod is reported in Fig 4A. Also in these plots, red lines fit experimental data, green lines represent simulations, while blue and red points are dawn and dusk respectively. Another defect concern the delay in the numerical results with respect to the experimental data. For instance in the 8h photoperiods (Fig 4B) there is a delay of 4h in the highest peak while in in the 18 photoperiods the delay is about of 6h. Anyway, the averaged dynamics well match simulation and biological results.

**Fig 4.**
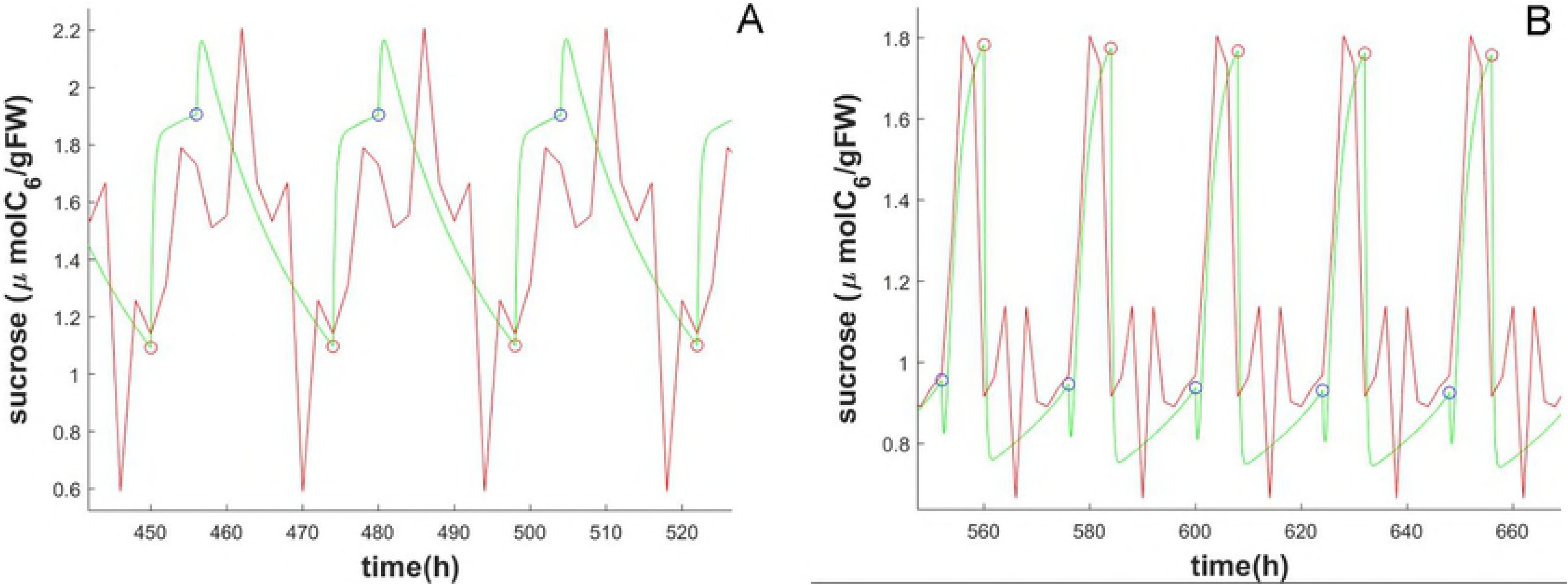
Behaviour inaccuracies. (A) Detail of sucrose content for 18h day length. (B) Detail of sucrose content for 8h day length.

The reason for such delay can be imputed to the *τ*_*as*_ function, which formulation require more specific investigations (see file S1 in supporting information). Both the delay and the smoother dynamic effects could imply an on/off control on *τ*_*as*_, with respect to some thresholds. Moreover, by assuming all parameters depending on photoperiods, it would be possible to further reduce the differences between model and biological data. However, we considered this validation sufficiently sharp (see next section 3.2), demonstrating the consistency of the model with respect to the literature.

Finally, we compared the relative growth rates (RGR). In [46], the RGR of fresh biomass of leaves is measured at the end of the night after 29 days. The mean values extracted from the literature are reported in Table 1 and compared with the RGR simulated by our model, obtaining an absolute error of 3.6283 · 10^−4^. To compute the RGR, we used the exponential formulation [73]:

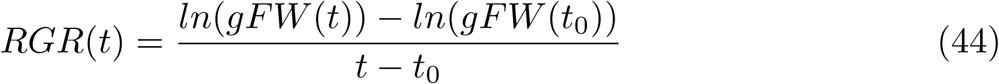

where t_0_ is the initial starting time.

**Table 1.** RGR obtained from literature and the model at different photoperiod.

| Source | 4h | 6h | 8h | 12h | 18h |
| --- | --- | --- | --- | --- | --- |
| [46] | 0.068 | 0.1135 | 0.1708 | 0.26 | 0.3065 |
| model | 0.0682 | 0.1135 | 0.1707 | 0.2600 | 0.3064 |

### 3.2 Model robustness

To evaluate the robustness of our model, we compared the numerical results with data extracted from several other papers. For instance, we tested the adaptation of the model to different light periods. In [42], it is shown how plant changes its metabolism and its sucrose content when day length changes from 8h to 16h (Fig 5).

**Fig 5.**
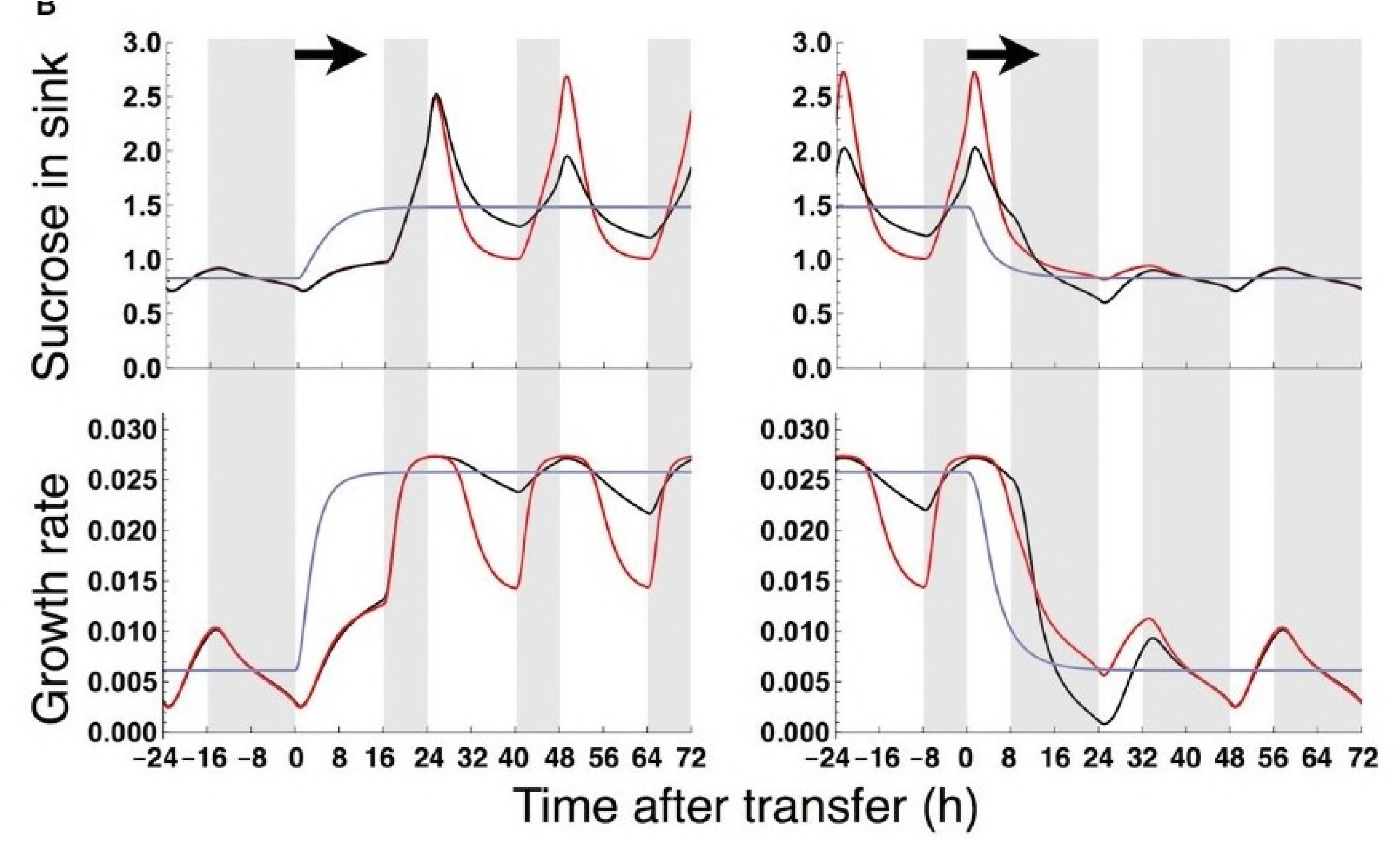
Changing behaviour in [42]. Dynamic of sucrose content in sinks in *Arabidopsis* when passing from 8h to 16h of light (in top left plot) and vice versa (in the top right plot). The black line refers to wild type plants.

Since, we are not explicitly considering sucrose content in sinks, we simulated only sucrose content dynamics passing from 8h light to 16h. The quantitative behaviour can not be the same, but the qualitative behaviour is supposed to be consistent. In fact, our model correctly predicts this dynamic as shown in Fig 6.

**Fig 6.**
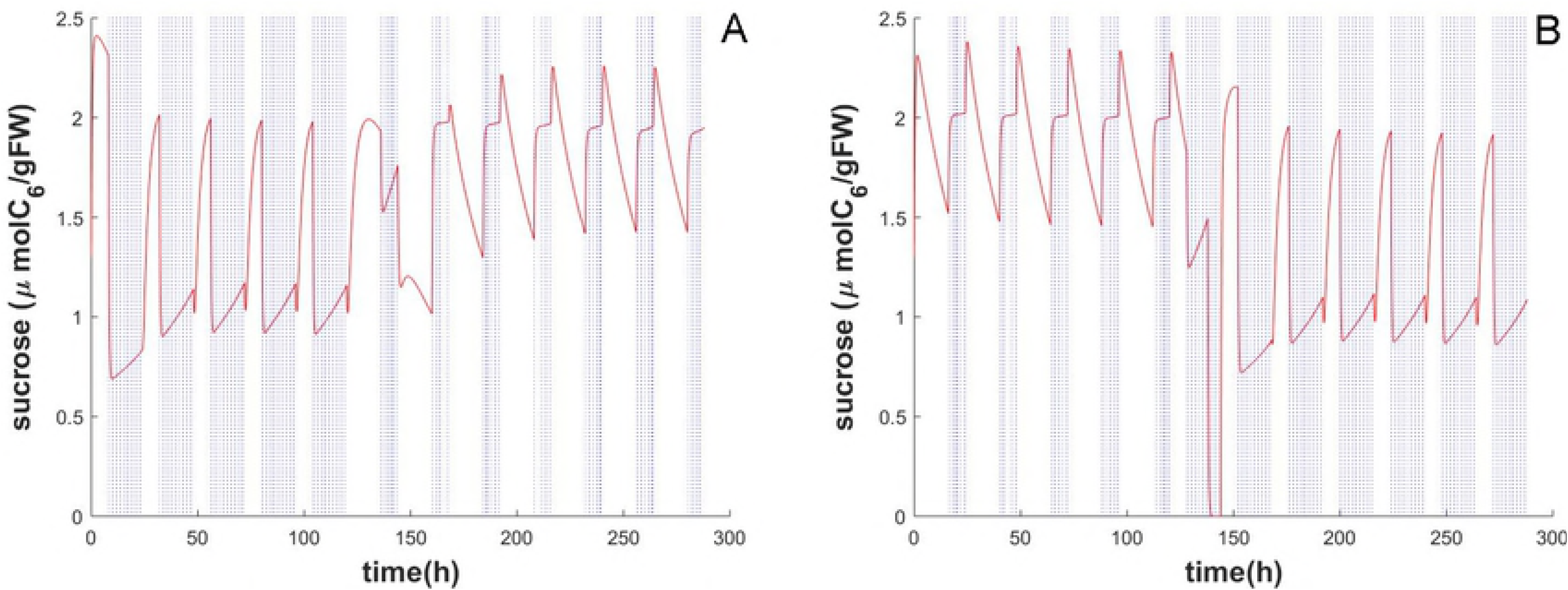
Changing behaviour in our model. (A) Our model simulation of sucrose content when passing from 8h to 16h of light. (B) Our model simulation of sucrose content when passing from 16h to 8h of light. Red line is the simulation, while blue dots represent dark periods.

Just like in [42], in the day when photoperiod changes, plant adapts its behaviour in an intermediate way, to fix it during successive days.

To verify the behaviour of *γ* we again referred to [42], in which *γ* varies along different photoperiods and it is estimated according to Table 2.

**Table 2.** Values of *γ* at different photoperiods as extracted from [42]

|  | 8h | 9h | 10h | 11h | 12h |
| --- | --- | --- | --- | --- | --- |
| $\gamma$ | 0.68 | 0.64 | 0.6 | 0.6 | 0.6 |

Nevertheless, [74] measured, in a standard experiment (12h of light), about 50% of starch accumulation. More recently, [75] verified changes in starch accumulation depending on photoperiod length and light intensity. The authors measured an accumulation range of [50%, 68.7%] for 6h photoperiods, and a range of [36%, 49.4%] for 12h photoperiods. The more light intensity, the less starch stored. To verify these data with our equations, we fixed non limiting initial conditions about nutrients and run the model for 10 days. Table 3 reports the averaged values of *γ*, which result to be consistent with data in the literature.

**Table 3.**
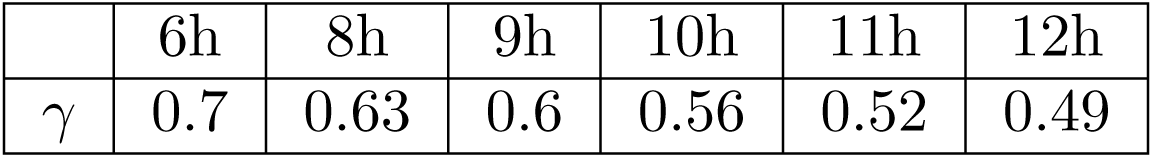
Numerical results for *γ* at different photoperiods.

|  | 6h | 8h | 9h | 10h | 11h | 12h |
| --- | --- | --- | --- | --- | --- | --- |
| $\gamma$ | 0.7 | 0.63 | 0.6 | 0.56 | 0.52 | 0.49 |

Another important validation is about the N:P ratio. In [23] it is shown how plant tends fast to optimal ratio when a missing nutrient is inserted in the environment where the roots grow. We fixed the value 10 according to the same reference and a light period of 12h. To test this behaviour, we fixed soil conditions and we varied the initial nutrient ratio. Starting from a very small and an higher initial ratio, the behaviour tends to approach the optimal ratio (Fig 7).

**Fig 7.**
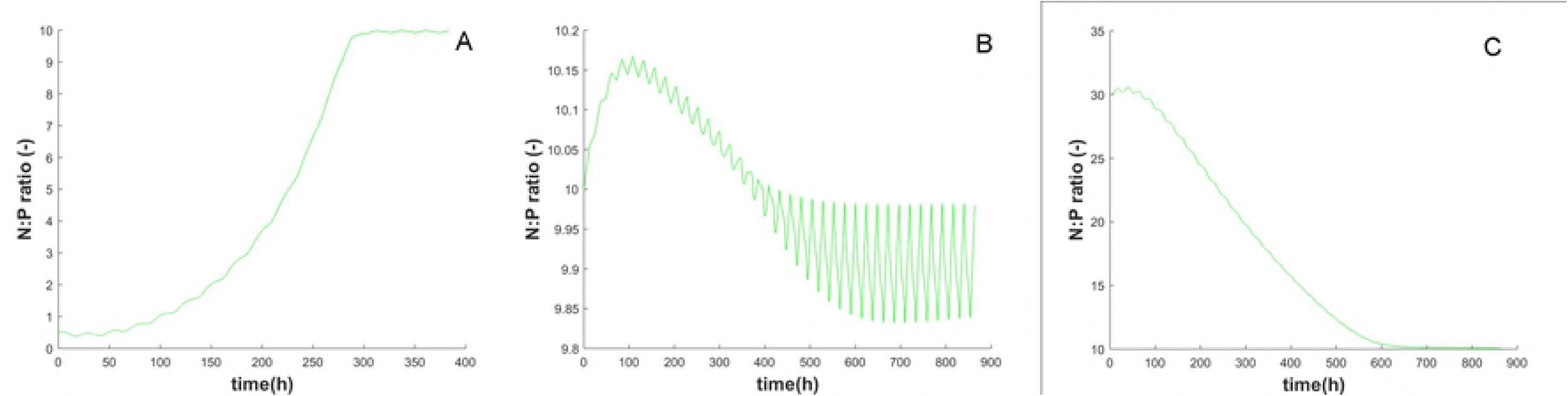
Evolution of N:P at different initial conditions. (A) N:P ratio in plant starting from a very little ratio (0.5); (B) N:P ratio in plant starting from the optimal situation; (C) N:P ratio in plant starting from a very large ratio (30).

Interestingly, [76] pointed out the role of N:P ratio in quantifying nutrient limitation on growth. Indeed, when we simulate different N and P soil contents, the plant reaches different stoichiometry ratios, as showed in Fig 8 (the point highlighted refers to N:P ratio in optimal nutrient soil content).

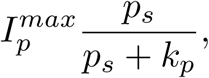

in poor *p* soils 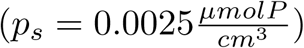 and rich *p* soils 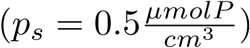. Then we run our model and computed hourly uptake:

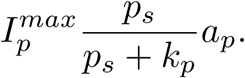

**Fig 8.**
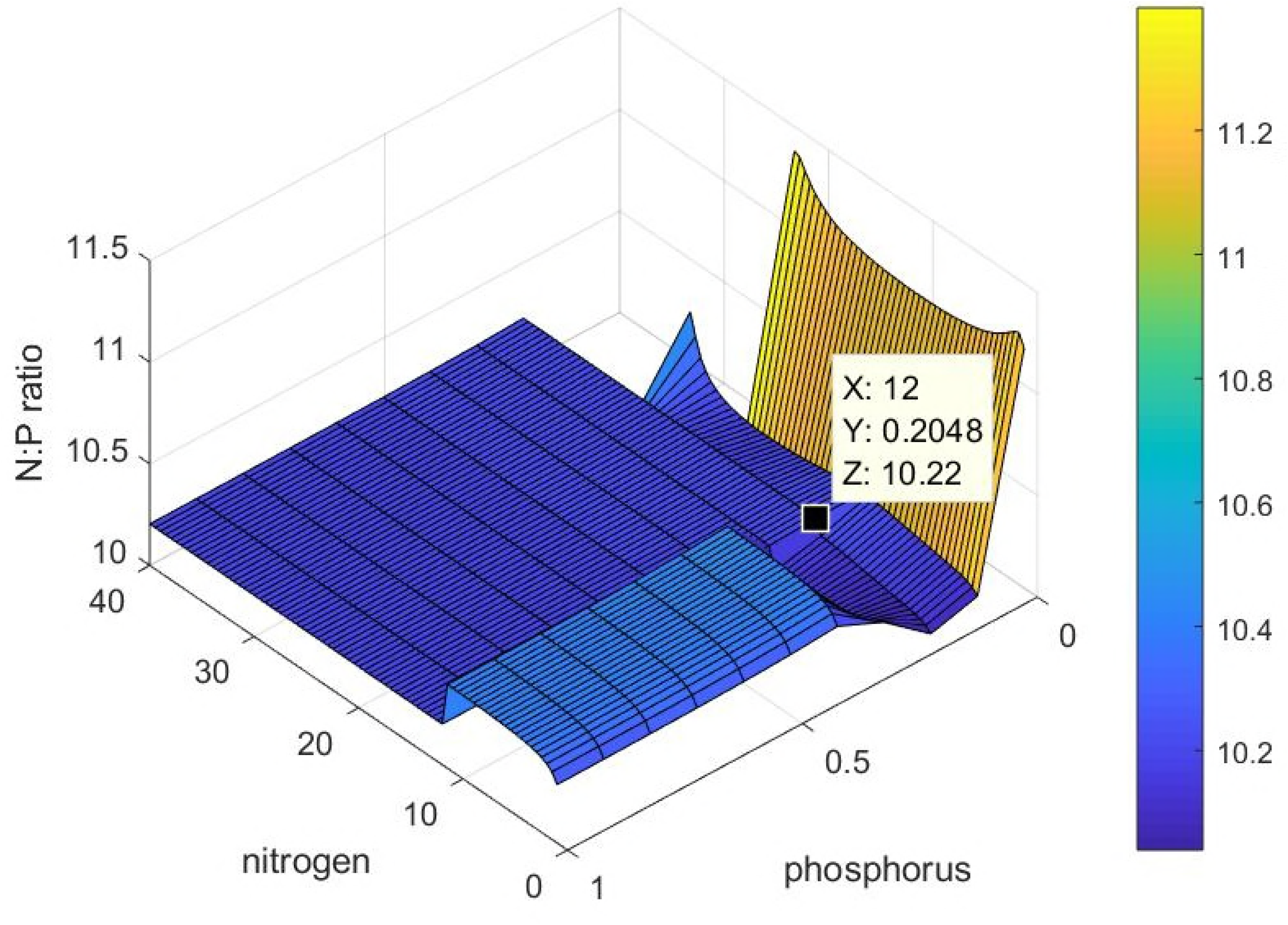
N:P dependence on soil nutrient contents. Moreover, when modelling the nutrient uptake, we assumed a dependence on the internal status instead of soil content. In [63], different values for both 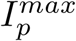 and *k*_*p*_ were proposed for different plants of *Arabidopsis*. We averaged among them in order to estimate the hourly phosphorus content:

In Table 4 there is the comparison between numerical results and our model, for 16h of light and 7 days of growth, showing a maximum error of 3.52%.

**Table 4.** Hourly uptake of P for poor and rich *p* soil, obtained from literature and our model, with relative error.

| Source | $p_s = 0.0025 \frac{\mu\text{molP}}{\text{cm}^3}$ | $p_s = 0.5 \frac{\mu\text{molP}}{\text{cm}^3}$ |
| --- | --- | --- |
| [63] | 0.108 | 0.142 |
| Model | 0.1075 | 0.147 |
| Relative error (%) | 0.46 | 3.52 |

Finally, we looked at the effects of excess and deficiency of P on growth. In [77] are reported two experiments on *Arabidopsis*, growing plants in 16h of light for 19 days under two different P treatments. The first treatment consists of a limited P medium 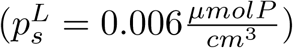, and the second consists of a saturated P medium 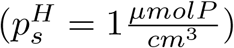. The authors measured the total dry biomass (DW) and the root:shoot ratio each 4 days. We used data from the 7^*th*^ day as initial point and we compared both fresh biomass and root fraction at day 11^*th*^, 15^*th*^ and 19^*th*^. Since our model is parametrized on fresh biomass, from [46] we estimated the relation DW= 0.08*FW*. Results, reported in Table 5, show an average error over the three periods of 0.028 ± 0.03 *gDW* in the rich soil and 0.00085 ± 0.00091 *gDW* in the poor soil, for the total biomass, and 0.43 ± 0.35 (%) in the rich soil and 1.71 ± 1.36 (%) in the poor soil, for the root fraction. The errors in biomass estimation, especially for the total biomass in rich soil, could be induced by the approximation adopted for FW to DW conversion. In fact, the water content could vary along the plant lifespan, affecting the relationship between dry and fresh weight.

**Table 5.** Total biomass and root:shoot biomass ratio obtained from literature and our model, with corresponding absolute error.

| Source | Parameter | Treatment | 11<sup>th</sup> day | 15<sup>th</sup> day | 19<sup>th</sup> day |
| --- | --- | --- | --- | --- | --- |
| [77] | Root fraction (%) | $p_s^H$ | 22.2 | 23.1 | 24.7 |
| Model | Root fraction (%) | $p_s^H$ | 22.63 | 23.96 | 24.7 |
|  | **Error** |  | **0.43** | **0.86** | **0** |
| [77] | Root fraction (%) | $p_s^L$ | 49.4 | 50 | 54.6 |
| Model | Root fraction (%) | $p_s^L$ | 45.96 | 51.59 | 54.7 |
|  | **Error** |  | **3.44** | **1.59** | **0.1** |
| [77] | total biomass (gDW) | $p_s^H$ | 0.01675 | 0.0398 | 0.057 |
| Model | total biomass (gDW) | $p_s^H$ | 0.01874 | 0.04979 | 0.1299 |
|  | **Error** |  | **0.00199** | **0.00999** | **0.0729** |
| [77] | total biomass (gDW) | $p_s^L$ | 0.00287 | 0.00425 | 0.004875 |
| Model | total biomass (gDW) | $p_s^L$ | 0.0025 | 0.0043 | 0.0070 |
|  | **Error** |  | **0.00037** | **0.00005** | **0.002125** |

The evolution in time of the biomass and the root:shoot biomass ratio obtained from our numerical simulation is also reported in Fig 9.

**Fig 9.**
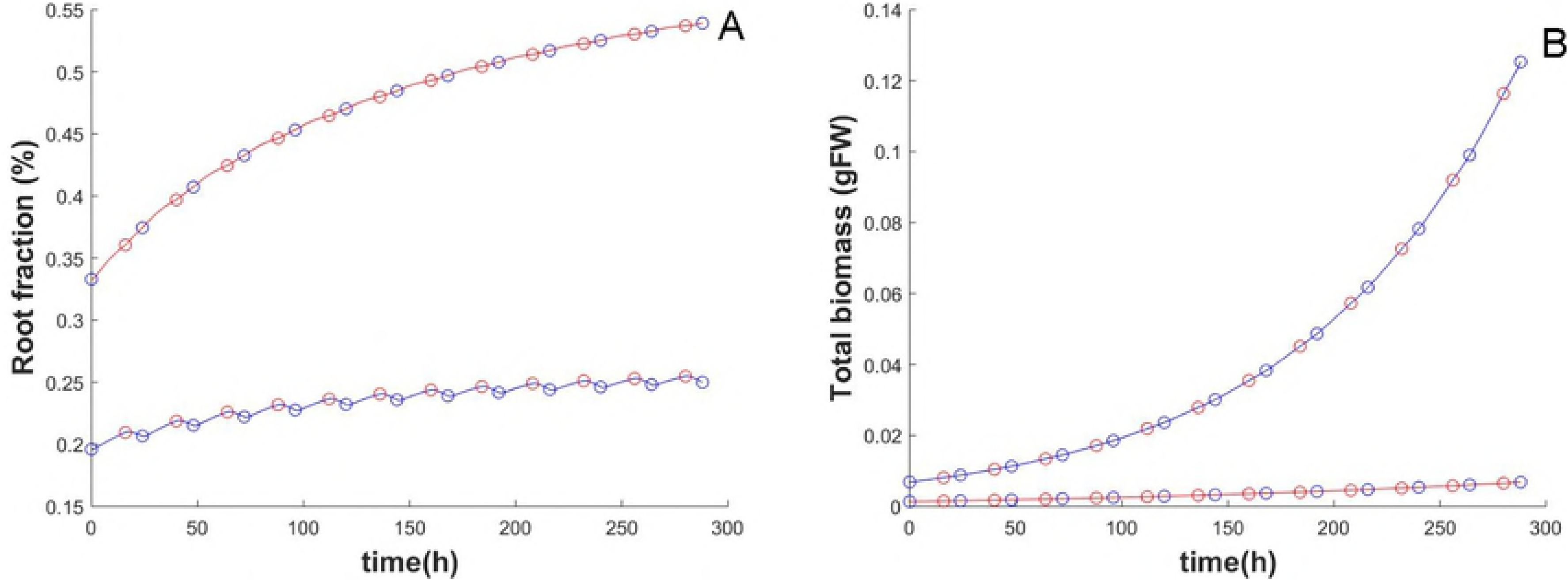
Validation test on [77]. (A) Root percentage under high and low phosphorus treatments. (B) Biomass dynamics under high and low phosphorus treatments. In blue, simulations for saturated P, while in red simulations for limiting P.

### 3.3 Root behaviour in heterogeneous soils

Up to now, soil conditions were stationary in time and nutrients were supposed to be only in the topsoil zone. In this section, we arbitrary fix 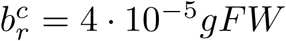 in order to study root system adaptation in heterogeneous soils. Results presented in the following refer to simulations of 20 days with 12h of light. We at first simulate root growth setting the topsoil rich in p, and the subsoil rich in n:

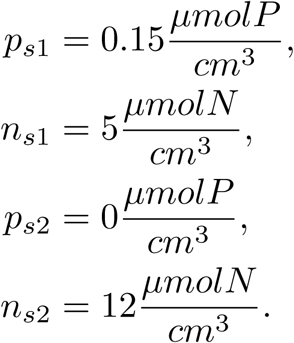

From Fig 10A we can observe that no *n* is experienced in the subsoil zone (and consequently no root is allocated) until the critical mass is reached. Then, plant starts allocating resources to elongate in the subsoil zone. Of course, at the beginning the more P reduces the N:P ratio that is immediately increased when the subsoil is reached and the more nitrogen is available (Fig 10B).

**Fig 10.**
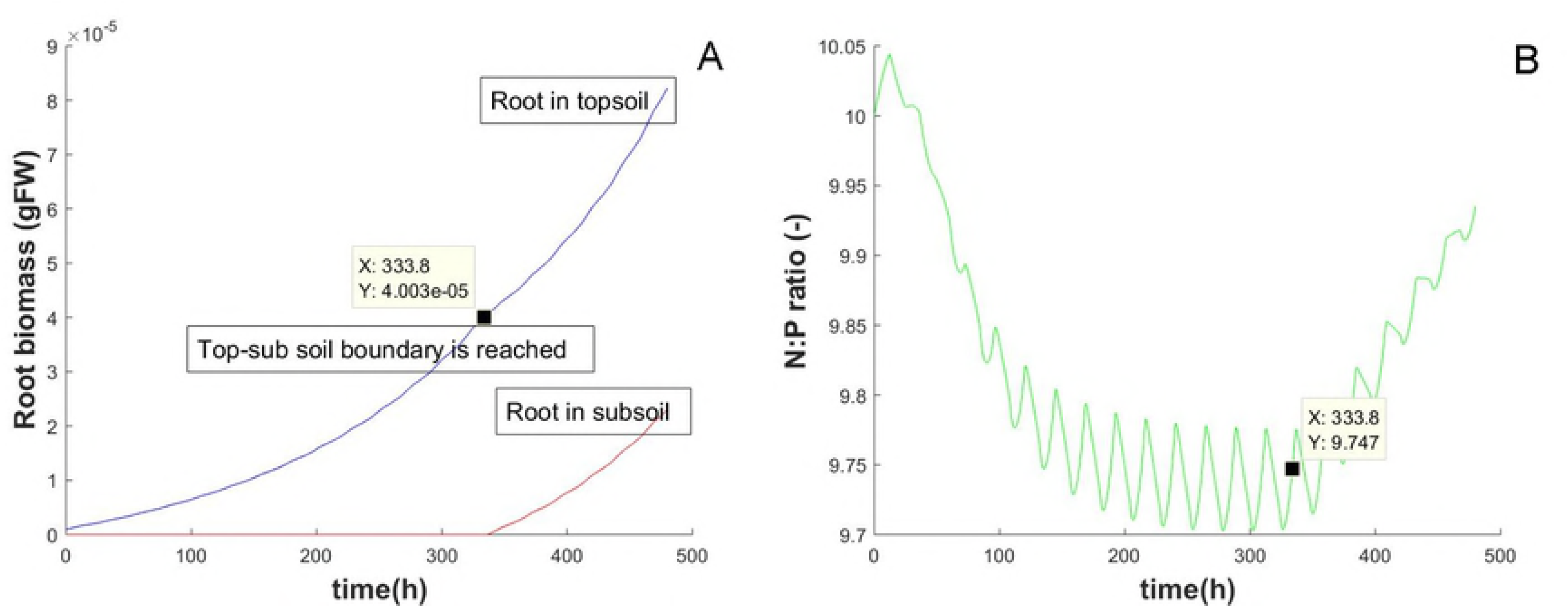
Two soil zones effects. (A) Root biomass growth in the two soil zones. Root growth in topsoil is drawn in blue line. Root growth in subsoil zone is drawn in red. (B) N:P ratio dynamics.

A more realistic simulation could be done assuming both nutrients initially placed in topsoil. But assuming that, day by day, nitrogen is transported by water from the topsoil to the subsoil:

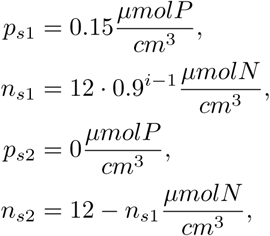

where the exponent i is the time (in days). Results are shown in Fig 11A. The less *n*_*s*1_ transported by water, the more *n*_*s*2_ accumulated in the subsoil.

**Fig 11.**
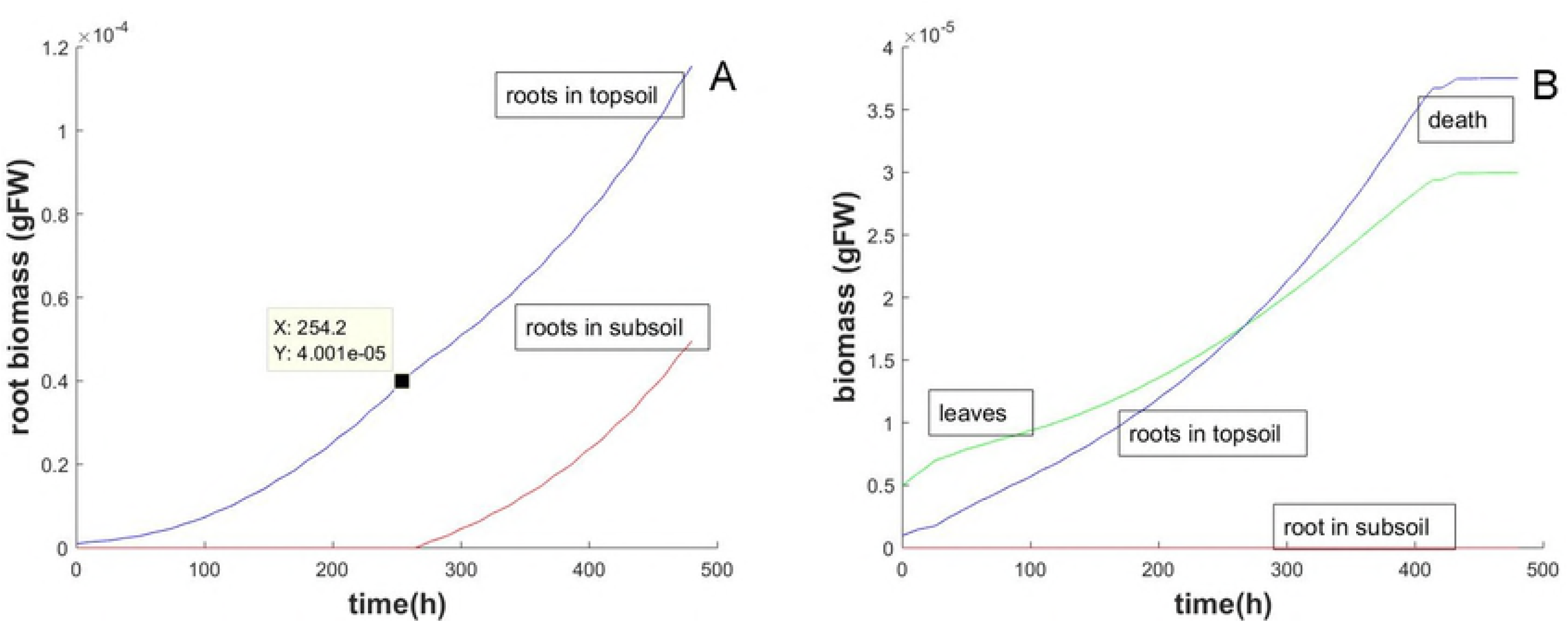
Example of plant death. (A) Root biomass growth in the two soil zones. Blue curve is the dynamic in the topsoil, red curve represents dynamics in subsoil. (B) Example of plant death. Blue line is the root growth in the topsoil, red line the root growth in the subsoil and green line is the aboveground growth.

To simulate a deeper nitrogen leak and then an higher topsoil zone missing n, the critical biomass *b*_*r*_^*c*^ can be increased. Moreover, it can happen the death of the plant, if nitrogen is lost faster than the time needed by roots for soil exploration. This condition can be obtained by the absence of growth. Let’s assume:

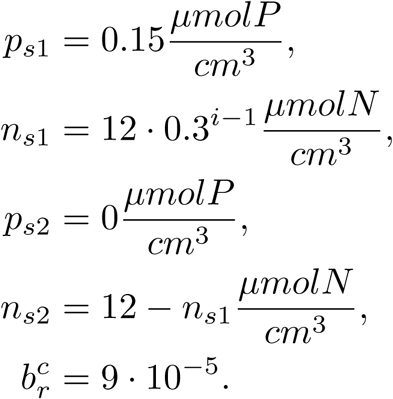

In this case, the root biomass is not able to grow up enough to reach the subsoil, although the greater investment in roots (different slopes in blue and green curves in Fig 11B). This way, the nitrogen is quickly exhausted and the plant is not able to survive.

We also verified the adaptive behaviour of plant in rich-soil zones (as reviewed in [55]):

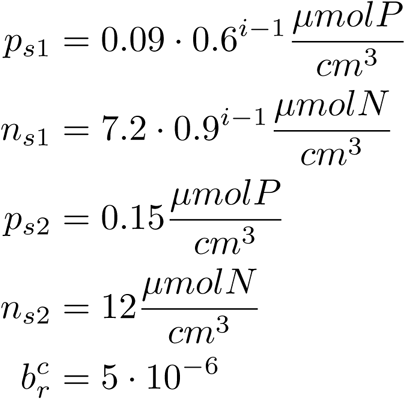

The plant used to grow in a more and more limiting topsoil, until the subsoil is reached. The more nutrient availability stimulates the growth in the deeper soil zone (Fig 12A).

**Fig 12.**
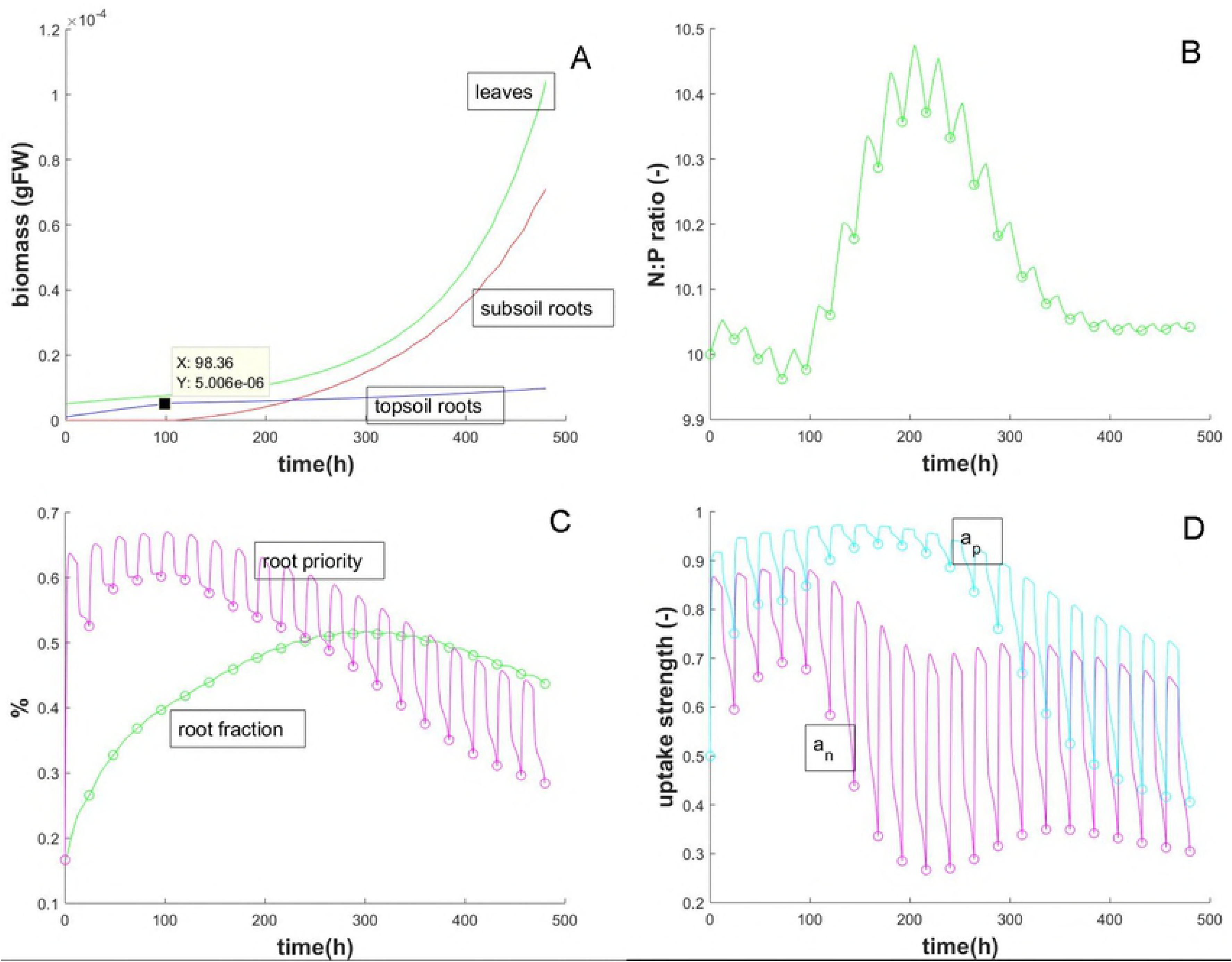
Changing soil zones. (A) Example of adaptation to different soil nutrient patches. The green line describes leaves, blue and red curve represent topsoil and subsoil roots respectively. (B) Example of N:P adaptation to different soil nutrient patches. (C) Example of the evolution of root fraction (green line) and root priority (*f*_*r*_) (magenta line) in adaptation to different soil nutrient patches. (D) Example of uptake adaptation to different soil nutrient patches.

Actually, the behaviour is more complex than what Fig 12A can express and the plant is, indeed, able to compute different signals at the same time. To show how our model simulate this complex network of stimuli, we provide additional results in the same framework of the previous simulation. Fig 12B shows the dynamics of N:P stoichiometry ratio and Fig 12C shows the evolution of root fraction and root priority (*f*_*r*_).

The adaptation of uptake signals *a*_*n*_ (magenta line for nitrogen) and *a*_*p*_ (light blue line for phosphorus) is instead presented in Fig 12D. In the end, the balance between uptake gain (*u*_*n*_ and *u*_*p*_) and nitrogen and phosphorus costs (*C*_*n*_ and *C*_*p*_) are shown in Fig 13.

**Fig 13.**
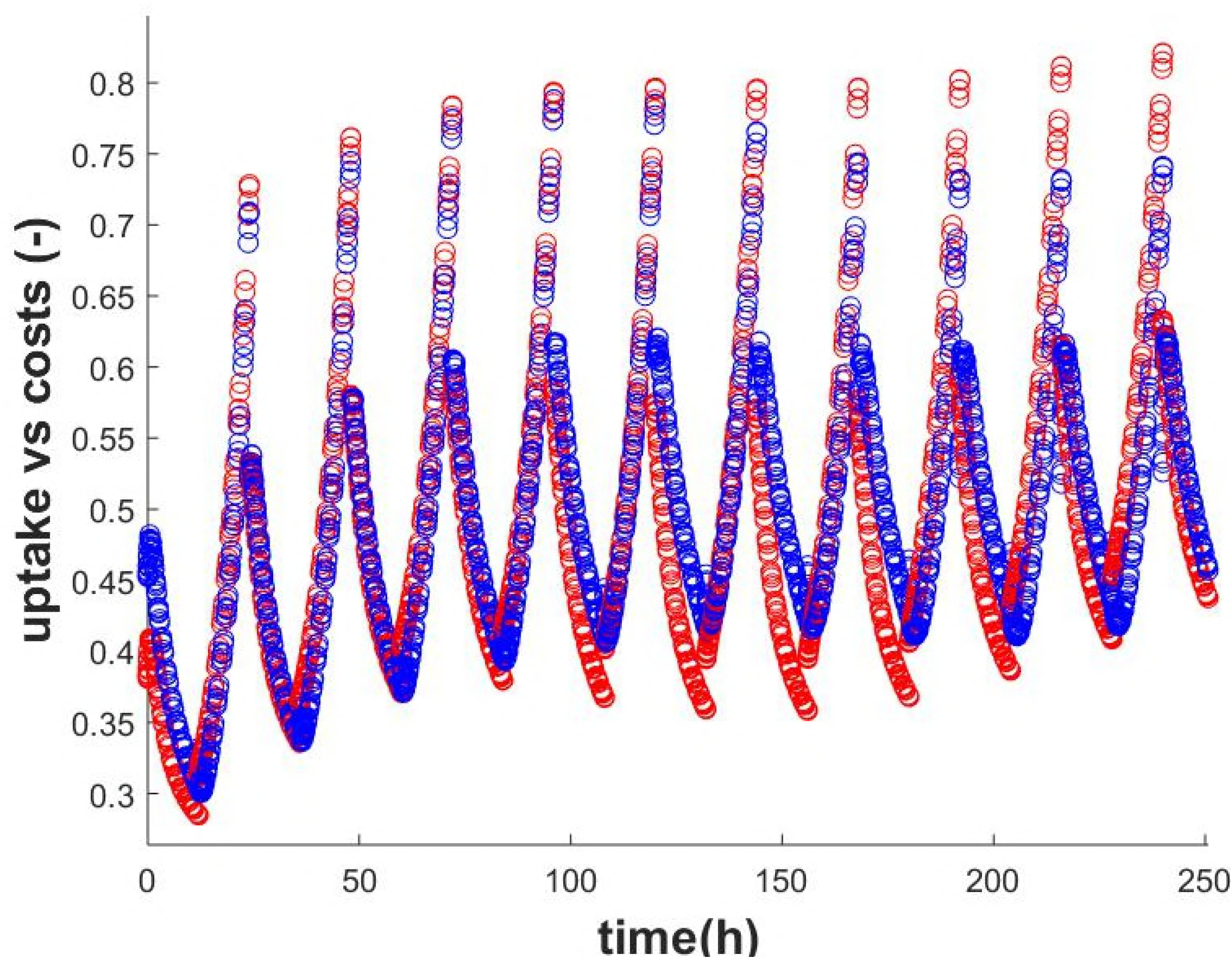
Example of adaptation to different soil nutrient patches. The blue dots represent nitrogen balance 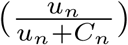, while the red dots the phosphorus one 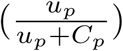.

All together, these plots allow us to appreciate a deeper simulation of plant adaptation. In fact, at the beginning, the plant experiences both low *n* and *p* in the topsoil. The limiting resources promote a greater uptake and a greater root priority. In Fig 12C the root fraction quickly increases. Moreover, the phosphorus is more limiting than nitrogen. In fact, in Fig 13 the averaged nitrogen balance is higher than the averaged phosphorus balance (the difference is evident at the beginning of the graph). This leads to a greater N:P ratio (Fig 12B) and a smaller *a*_*n*_.

However, according to [55], differences in uptake costs and root growth costs define the nutrient affinity improvement of plant or the increasing of root fraction. In the phosphorus limiting case, being the soil exploration useless due to nutrient deficiency (until the subsoil is reached), it is more convenient to increase phosphorus affinity *a*_*p*_ more than root fraction (light blue curve in Fig 12D and magenta curve in Fig 12C). On the other hand, when the rich soil is reached, plant cannot immediately change its strategies due to the concept of memory [18]. In fact, plots show no changes in dynamics after having passed the topsoil boundary. Anyway, more phosphorus is now available and the unmodified strategy leads to luxury uptake of phosphorus and a critical reduced N:P ratio (Fig 12B). To stabilise the stoichiometry ratio, plant starts reducing phosphorus affinity. Moreover, in rich soil zones, the uptake is more convenient than root proliferation, so that root priority is less strong than *a*_*n*_. In the end, when both the optimal ratio is reached and the memory is coordinated with the new soil nutrient content, plant can stabilise its signals.

### 3.4 Leaf-root competition

We simulate leaf and root competition by fixing parameters 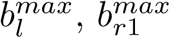 and 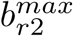. For this analysis, we assumed only one soil zone with optimal nutrient contents and we performed a simulation of 20 days. When assuming 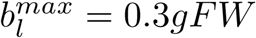, the critical leaf biomass is reached after *t* = 343h. However, even after the critical biomass, plant does not stop growing (Fig 14)but the sucrose is not enough to sustain the actual growth (Fig 15A). This is the signal that later will reduce the growth. Both leaves and roots decrease with respect to a plant not experiencing the competition (Fig 15B and 15C). Of course, there are no reasons why affinity should change (for example in Fig 15E we report the phosphorus affinity *a*_*p*_ being similar also the *a*_*n*_ dynamics). Letting conditions unchanged and simulating 50 days, a new equilibrium is reached by the plant (Fig 15E).

**Fig 14.**
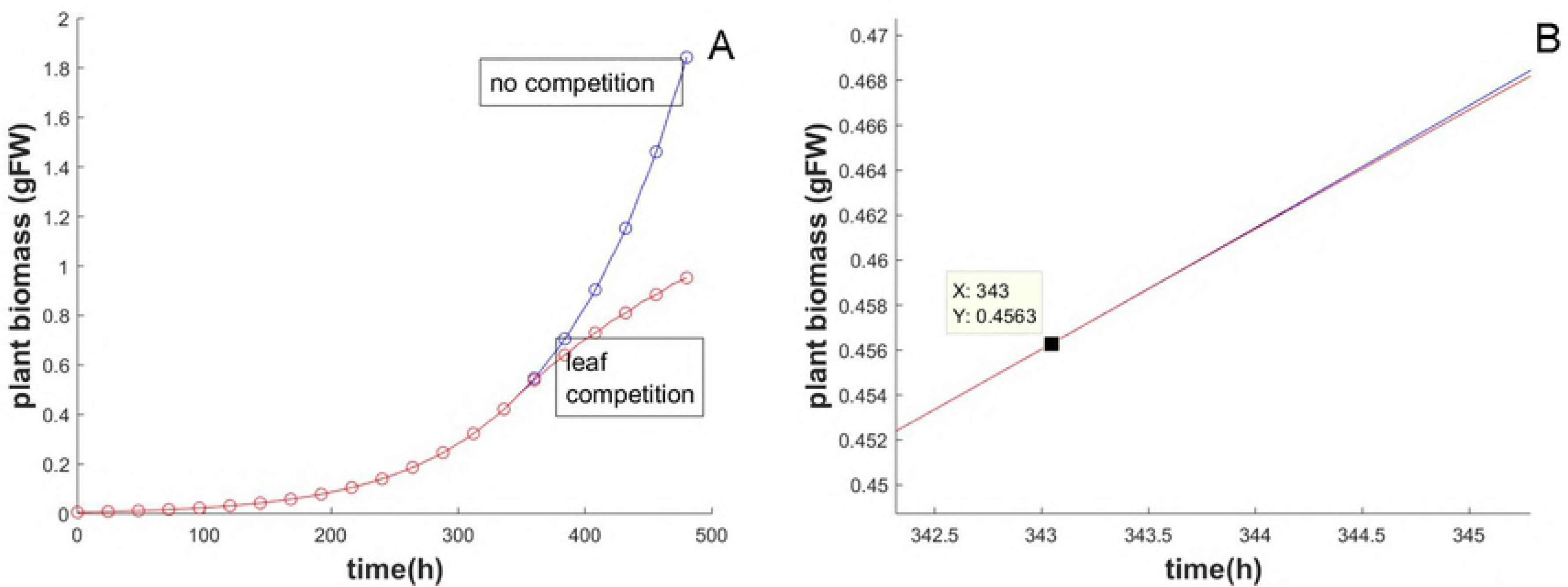
Effects of leaf competition on plant biomass. (A) Total behaviour. (B) Detail close to = *t*324.4h. In blue the case of no competition, while in red the case of 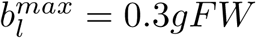.

**Fig 15.**
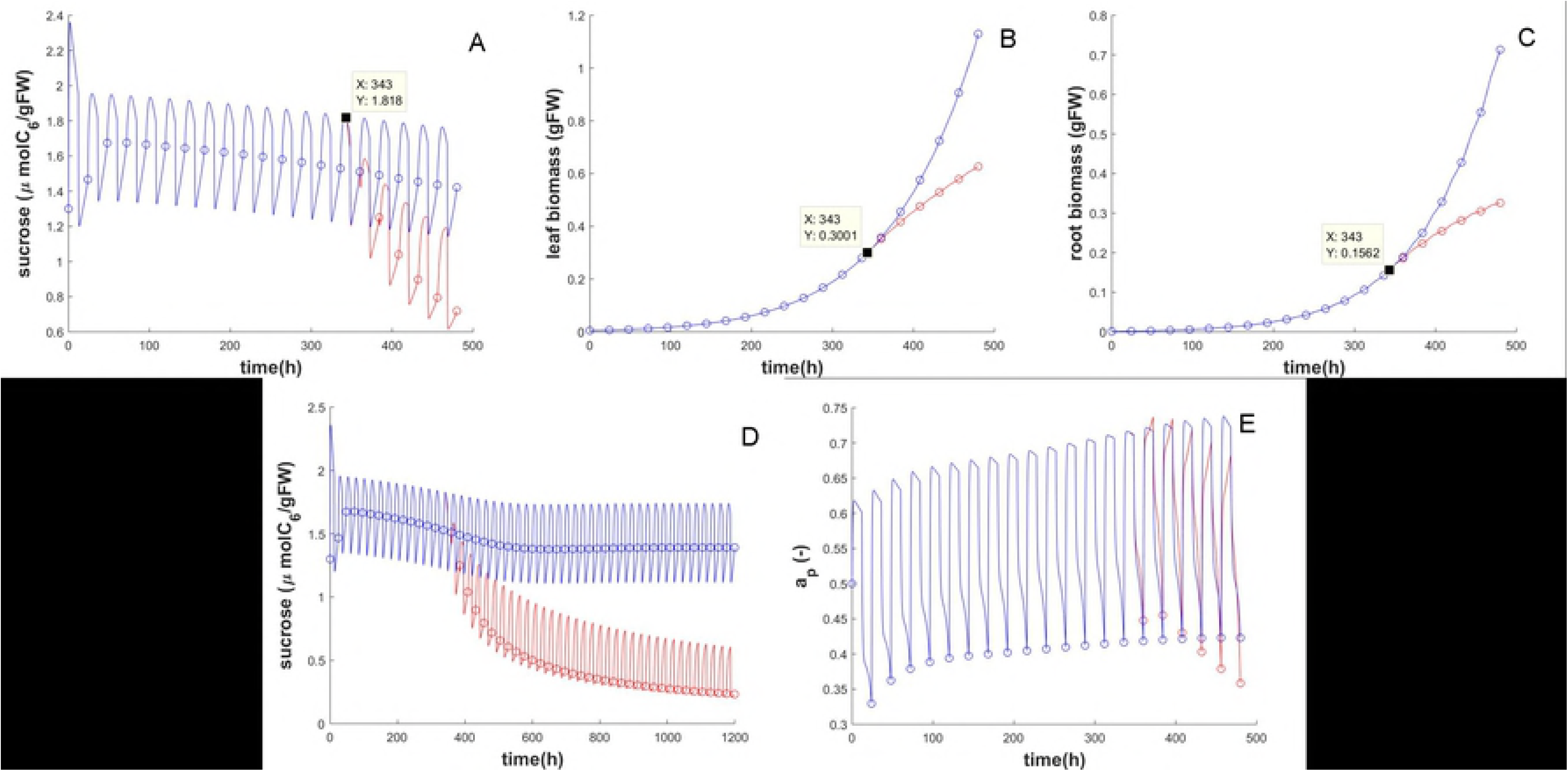
Effects of leaf competition on plant biomass. In blue the case of no competition, while in red the case of 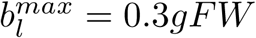. A) Sucrose dynamics (B) Leaf dynamics (C) Root dynamics (D) Sucrose dynamics for a dynamic of 50 days showing a new stabilisation (E) Adaptation of phosphorus uptake feedback (being the nitrogen one similar)

### 3.5 Root overproduction

According to [78] and [79], the competition among adjacent roots occurs when the respective rhizosphere overlaps the other one, reducing the uptake for unit of root mass. This is consistent with our modelling approach (see section 2.6.1). For this evaluation, we assumed only one soil zone, 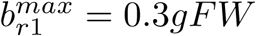 and, initially, no leaf competition. We performed simulations for 20 days at 12h of light, and the results are shown in Fig 16. As expected, plant promotes leaf growth instead of the root one.

**Fig 16.**
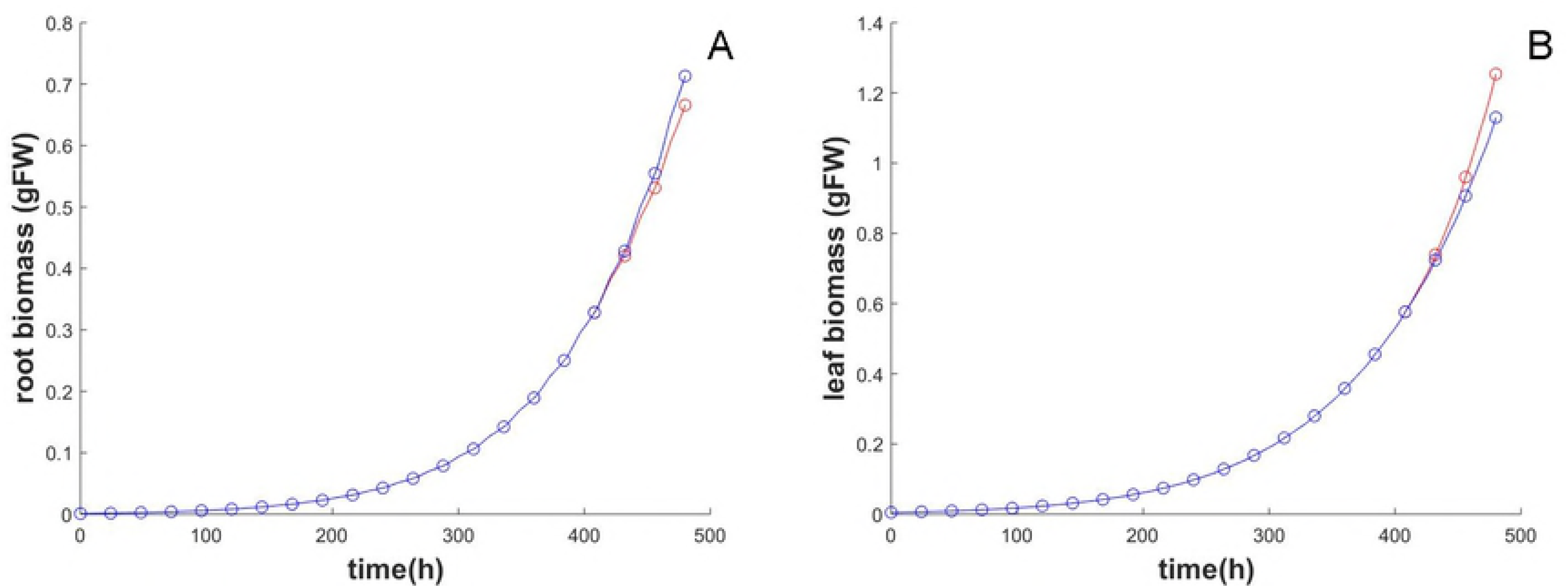
Effects of root competition on roots and leaves. The blue curve is growing without competition.

However, for a more realistic framework we should take into account also leaf competition. For this reason, we further simulate the behaviour in two conditions: 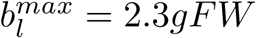 and 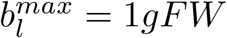.

In the former case (greater leaf critical biomass), the strategy is dangerous because favouring leaves than roots, roots can not absorb enough nutrients to sustain the faster leaf growth. At the beginning, the total biomass is greater with respect to the no competition framework (red curve in Fig 17A).

**Fig 17.**
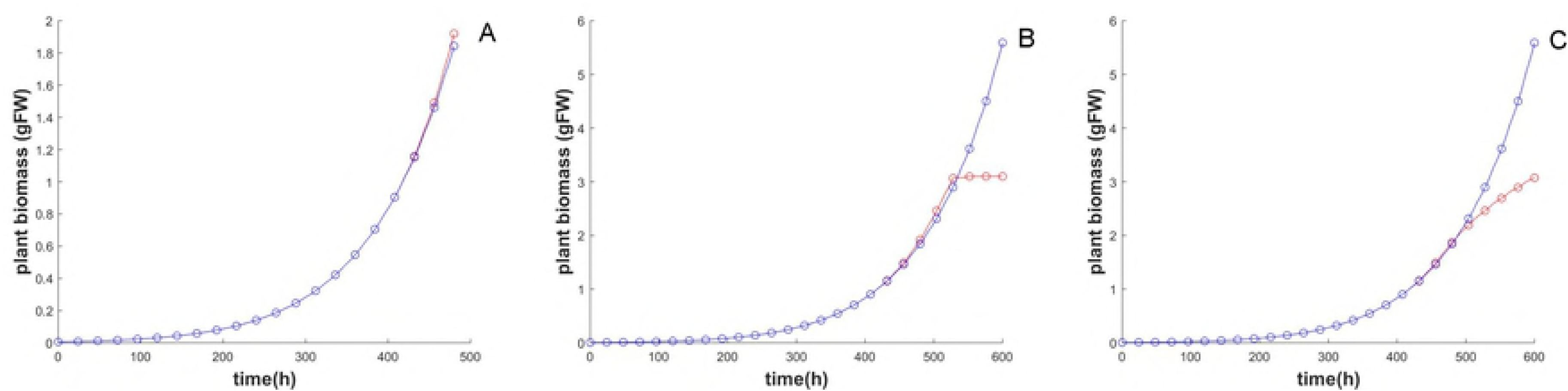
Total competition effects. (A) Effects of later leaf competition and faster root competition on plant biomass. (B) Effects of plant death due to a later leaf competition. (C) Effects of faster leaf and root competition in plant biomass.

After few days (25 instead of 20 days) the plant dies (red curve in Fig 17B). In the second case, the leaf competition happens when the plant can still sustain its development. The growth still hold but in a slower way and the plant does not die (red curve in Fig 17C).

Finally, let’s assume two optimal soil zones 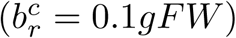 in a competing framework (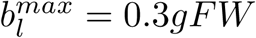 and 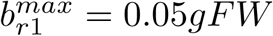) for 30 days. Initially, the plant experiences the competition inducing an unbalance between nutrients: the N:P ratio is not anymore optimal (Fig 18A). After having reached the subsoil, the plant promotes the root growth in the subsoil (Fig 18B and 18C) and stabilises again the optimal ratio.

**Fig 18.**
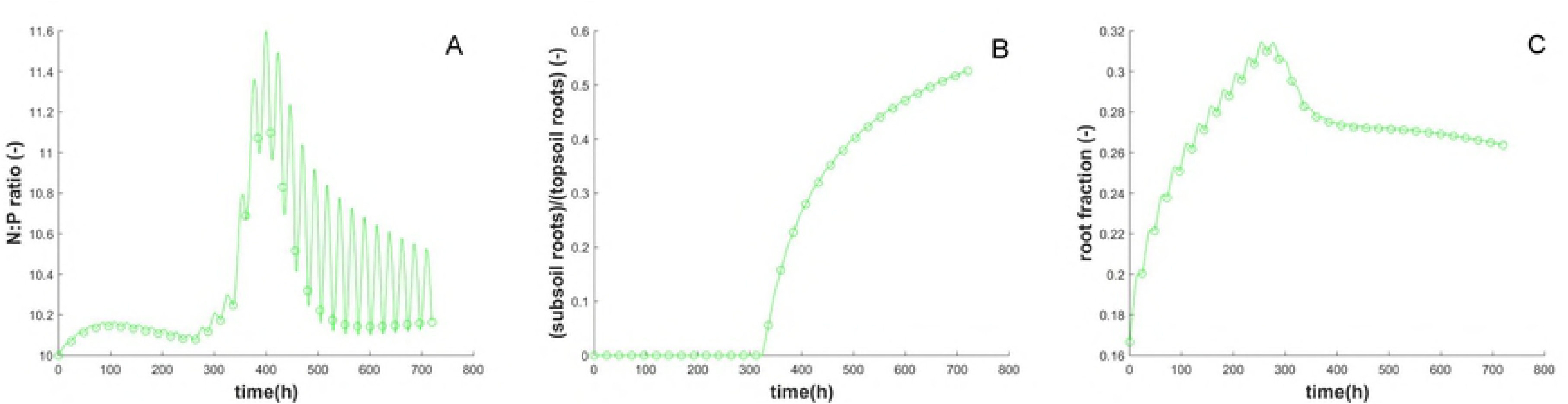
Effects of leaf and root competition in two soil zones. (A) N:P ratio. (B) Subsoil roots vs topsoil roots. (C) Root to shoot ratio.

## 4 Discussions

In this paper, we took stock of existing biological literature extrapolating the main evidences of intra-plant communication in the form of interactions between the dynamics of internal processes in plants. By the experimental data obtained from literature, it was possible to point out the relationships between such processes and to define realistic parameters for the equations proposed in the model.

To date, the mathematical models of plant growth actually disregard plant communication. They mainly focus on the description of a single and specific process such as photosynthesis, uptake or water transport; they are devoted to the description of the chemical interaction among hormones; or limited to a single nutrient.

To have a glance on the internal communication driving the growth, and differently from the state of the art, we proposed here a model that integrates at the same time several processes (photosynthesis, starch degradation, nutrients uptake and management, biomass allocation, maintenance), different signals (circadian clock and growth stimulus) and nutrients (sucrose, nitrogen and phosphorus).

The overall system extends results from previous models ([20, 41]) and proposes an approach, that could be addressed to biologists, for analysing interactions not yet investigated. For example, thanks to the model it is possible to evaluate how photosynthesis intensity and sucrose production rate are reduced or increased according to different stoichiometry ratio values, or which process is mainly, and earlier, affected by nightly starvation. By evaluating the dynamics of the priority signal *f*_*r*_ when the environmental conditions change, it is possible to investigate on the effects of starch accumulation during biomass allocation. Furthermore, one may inquire responses to leaves and roots overproduction or photosynthesis limitations due to light and stored starch.

Also, new questions about the nutrient uptake mechanism emerged from the model. In fact, the classic Michaelis-Menten dynamics ([63–65]) was generalised to include internal stimuli. The accuracy of results suggests the need for new biological experiments to additionally investigate on this mechanism.

Comparing our model with [20, 42], a new relation has been defined for the growth, which does not depend anymore just on nutrients availability, but it takes into account also starvation and efficiency. Here, by efficiency of an organ (which could be a leaf or a root), we mean the ability of that organ to provide the amount of resources needed for the correct functioning and growth. For example, the roots efficiency is the ability to provide both nitrogen and phosphorus such that nutrient levels are enough to sustain hourly costs of nitrogen (*C*_*n*_) and phosphorus (*C*_*p*_). On the other hand, leaves efficiency is the ability to produce enough sucrose and starch so that all respiration costs 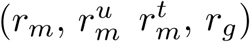 are covered by the available sucrose. Hence, as reviewed in this paper, plant tries to never reach minimum nutrients thresholds, while keeping monitored level of stores to avoid consuming energy unnecessarily.

What the model can show, by analysing these dynamics, is a dense network of signals and feedback that enforce the concept of intra-plant communication allowing a deeper understanding of how this internal network is connected.

Nevertheless, the model can be used to better define biological concepts like *performance*, nutrient *efficiency* and *sink priority*.

In [24], *performance* is defined as *”the ability to acquire resources and survive in the presence of competition or in stressful environments where physiological limits are reached”*. The problem in measuring the performance refers to the problem of defining the parameter to be checked in stressful conditions: either the biomass, sucrose production, nutrient uptake or photosynthate allocation are important parameters that can provide an idea about the ability of the plant to survive. Our model allows to easily estimate the plant performance by simulating limiting scenarios and measuring sucrose, nitrogen and phosphorus levels with respect to the critical thresholds.

In [80], the *efficiency* in using nutrients is generally defined as: the *”measure of how well plants use the available mineral nutrients*. *It can be defined as yield (biomass) per unit input (fertiliser*, *nutrient content)”*. A similar definition can be found in [25] for efficiency in the use of water, and related to photosynthesis efficiency.

Clearly, the biomass parameter is not enough to measure the efficient use of nutrient; in fact, just considering this parameter, a simple imbalance in roots and leaves biomass could imply a not efficient use of resources, whereas it is not always the case [81]. In our model, the efficiency emerges from relationships among uptake and nutrient costs (from Eq (29) and (30)). In fact, it is possible to note a nutrient efficiency in the first ratio, relating uptake (*u*_*n*_ and *u*_*p*_) with costs (*C*_*n*_ and *C*_*p*_), and a photosynthesis efficiency in the last ratio, which relates uptake and photosynthesis. In addition, Eq (13), relating sucrose and its critical thresholds, gives an idea about sucrose efficiency.

Finally, [82] defines the strength of sinks, or *sink priority*, as *”the competitive ability of a sink organ to import photoassimilates*, *and depends on both its physical (size) and physiological (activity) capabilities”*. While the size is a clear parameter and it can be easily related to the biomass of the organ, defining the activity of a sink is more difficult. In [83], the term *sink activity* is referred to all processes involved in using resources, from the synthesis of new tissues to the maintenance of the already existing ones. This activity can be quantified as the net assimilation of resources in tissues minus losses due to respiration and exudation. However, in [84] it is noted how the previous definition of sink strength is not enough to explain how sucrose is allocated in the plant. In fact, processes affecting osmotic concentrations in sinks need to be taken into account for resources allocation. Furthermore, in [83], is highlighted the importance of associating the source-sink interaction with the whole plant growth, even if no hint about how to do it is given.

In our model, both the sucrose partitioning signal (*f*_*r*_) and the root competition function (*e*_1_) are used to describe this *sink priority*. They depend on resources content (taking into account osmotic concentration) and, indirectly, on the biomass (size) and internal processes (activity). In this way, it is possible to connect the general definition of sink strength with the concept of osmotic concentration, other than proposing a practical association between plant growth and source-sink interactions, through the effects of *f*_*r*_ and *e*_1_ on the biomass dynamics (leaves *b*_*l*_ and roots *b*_*r*_).

## 5 Conclusions

Plants are an important model of adaptation by acting with morphological and physiological responses to environmental changes. The richness in internal and external signals that plants use for regulating their growth strategies are opening new frontiers about the ability of plant to communicate ([1]). The understanding of the processes and actors behind their communication is of fundamental importance not only for improving the knowledge of plant functioning, but also for helping in translating such abilities in many other fields for different applications (e.g. for collaborative tasks in swarm robotics [23], for resolution of combinatorial problems [85]).

The mechanisms of plant communication are being studied through self-recognition cues, underground interactions with fungi or other plants, volatile organic compounds, electric and chemical signals ([86–88]). In this paper, we investigated in the intra-plant communication by analysing the possible cues activating the adaptive growth responses in a single plant and describing the dynamics of such internal processes and signals with a mathematical model. Our assumptions and formulations have been deeply analysed in comparison with other models from the state of the art, highlighting the adopted extensions and improved potentialities.

Main difference with other models is having included several different elements (e.g. photosynthesis, starch degradation, multiple nutrients uptake and management, biomass allocation, maintenance) and their dynamics all at once considering their interactions and effects on the growth, thus showing the potential intra-communication in plants; whereas, strength of our model lies in having based assumptions and formulations on biological evidences and laboratory experiments collected from the state of the art. This approach allowed to evaluate several parameters from the papers thus reducing the number of free variables, and at the same time, to maintain a high level of details with good agreements with the biological model.

To test the robustness of our model, we compared the biomass, sucrose dynamics, photosynthesis partitioning rate, uptake strength and sinks priorities not using custom experiments, but encouraging the comparison with different independent published biological data. Indeed, we validated the model by comparing experimental results with results obtained in our simulations, reproducing conditions of growth similar to the biological counterpart. All validation tests show high accuracy in results and very small errors (e.g., we obtained 3.52% as maximum relative error in nutrient uptake comparisons (Table 4) and 3.44% as maximum percentage error for biomass estimation in low phosphorus soil tests (Table 5)); where, the errors obtained are likely induced by divergences of initial conditions between biological and numerical experiments. However, both the qualitative behaviour and the quantitative values agree with data in literature.

Moreover, we simulated different growth scenarios. Interestingly, in stress conditions, e.g. when the saturating thresholds were reached, the dynamics showed a fast adaptation of the plant with a self-optimisation of internal resources and feedback stimuli, without the need of a *a-priori* optimisation assumption. The same self-adaptive behaviour was evident when branching vs elongation was analysed. This optimising behaviour is realistic but difficult to be biologically demonstrated. However, differently to some models [4, 19] that directly start by assuming the existence of an internal optimisation function which drives plant dynamics, in here we demonstrated it to be a consequently behaviour emerging from the dynamics of internal processes. Therefore, our model enforces the idea of an emerging behaviour in plants making use of an internal communication network for optimising the use of the internal resources, the exploration of the soil and the adaptation of internal processes, and results in an efficient growth strategy [23].

Beside important biological implications and analysis, this model can also leverage the development of plant-inspired algorithms for the control of exploratory robots ([89], [23]). For instance, the equations, identically replicated on each robot, can describe the behaviour of the robot and represent the ability of robotic roots to colonise a certain area (plant branching) or to growth further in exploration (plant elongation), according to the internal signals (into the colony of robots) and the measurements from the environment.

Future steps will include from one side, the implementation of the proposed behaviour in robotic solutions, while from a biological perspective several improvements can be adopted. For instance, the saturating thresholds can be replaced by death terms that need to be estimated via laboratory experiments. Stem and fruit dynamics can be introduced for a more complete description. Moreover, the concepts for resources allocation, analysed in this paper only in roots, could be used to study resources partitioning in leaves. Additionally, from a mathematical point of view, partial differential equations can replace root dynamics to analyse spatial diffusion of roots. Moreover, the model is easily embeddable with more detailed formulations for photosynthesis, light, temperature, starch degradation, respiration or exudation in order to make the model more and more detailed.

Ultimately, the model can become a powerful tool for the study and validation of biological hypothesis; step by step introducing additional dynamics, it can describe complex behaviours and help in finding the signals driving them; it can further be integrated for study the interactions among two or more plants or for instance, to highlight the effects of having invasive or not-invasive species. If coupled with nutrients dynamics in soil, the model can provide useful tools for optimising outer resources distribution to reach a target biomass. As well as, a coupling with a stochastic model of temperature and light can allow a study of a seasonal adaptation of the plant.

## 6 Supporting information

**S1 Appendix. Starch degradation comparison.** A more detailed starch degradation function is integrated in the model. The results are discussed and new open questions are proposed about this complex chemical process.

